# Nasal commensals reduce *Staphylococcus aureus* proliferation by restricting siderophore availability

**DOI:** 10.1101/2024.02.20.581135

**Authors:** Y Zhao, A Bitzer, JJ Power, D Belikova, B Torres-Salazar, LA Adolf, DL Gerlach, B Krismer, S Heilbronner

## Abstract

**Background:** The human microbiome is critically associated with human health and disease. One aspect of this is that antibiotic-resistant opportunistic bacterial pathogens such as methicillin-resistant *Staphylococcus aureus* can reside within the nasal microbiota which increases the risk of infections. Epidemiological studies of the nasal microbiome have revealed positive and negative correlations between non-pathogenic species and *S. aureus*, but the underlying molecular mechanisms remain poorly understood. The nasal cavity is iron-limited and bacteria are known to produce iron-scavenging siderophores to proliferate in such environments. Siderophores are public goods that can be consumed by all members of a bacterial community. Accordingly, siderophores are known to mediate bacterial competition and collaboration but their role in the nasal microbiome is unknown.

**Results:** Here we show that siderophore acquisition is crucial for *S. aureus* nasal colonization *in vivo*. We screened 94 nasal bacterial strains from seven genera for their capacity to produce siderophores as well as to consume the siderophores produced by *S. aureus.* We found that 80% of the strains engaged in siderophore-mediated interactions with *S. aureus.* Non-pathogenic corynebacterial species were found to be prominent consumers of *S. aureus* siderophores. In co-culture experiments, consumption of siderophores by competitors reduced *S. aureus* growth in an iron-dependent fashion.

**Conclusions:** Our data show a wide network of siderophore mediated interactions between the species of the human nasal microbiome and provide mechanistic evidence for inter-species competition and collaboration impacting pathogen proliferation. This opens avenues for designing nasal probiotics to displace *S. aureus* from the nasal cavity of humans.

## Background

The human body is colonized by a multitude of bacteria from different genera that is collectively called the microbiome. The human microbiome is fundamentally associated with human health and disease. One aspect of this is that Antibiotic-Resistant Bacterial Pathogens (ARBPs) can hide within microbiomes of healthy individuals and can cause invasive disease if, for example, they are transferred into surgical wounds after invasive interventions within hospitals (1). *Staphylococcus aureus* is a prime example in this regard. The pathogen colonizes the anterior nares of approximately one third of the human population and colonization is a major risk factor for infection (2,3). *S. aureus* infections cause severe morbidity and mortality and are frequently difficult to treat due antibiotic-resistant lineages such as methicillin resistant *S. aureus* (MRSA). We know surprisingly little about the factors that determine whether an individual can be colonized by *S. aureus.* Host genetics and environmental conditions have only a moderate influence on *S. aureus* colonisation (4,5). It is increasingly recognized that the presence of certain commensal species is important. Epidemiological analyses of nasal microbiomes showed the presence of *Finegoldia magna*, *Dolosigranulum pigrum* and *Simonsiella* spp., to be negatively correlated with the presence of *S. aureus,* and corynebacteria are associated with a reduced absolute number of *S. aureus* cells (5,6). However, few studies have investigated the molecular interactions between *S. aureus* and other nasal commensals that underly these observations. Production of antibacterial molecules is known to shape the composition of microbial communities (7) and has been shown to be important for the displacement of *S. aureus* by certain commensals (8–10). In nutritionally limited environments, competitive or collaborative exploitation of natural resources is important in shaping microbial community structures (11,12). However, little is known about this in the context of the nasal microbiome (13). A well described mechanism of cooperation between bacterial cells is the secretion of “public goods” that are costly to produce but can be used by the entire community (14). Iron-scavenging siderophores (SPs) are a classical example. SPs are produced by many bacterial species and allow iron-restriction in many environmental and host-associated habitats to be overcome. Due to their costly production, organisms called “cheats” that consume SPs produced by others but lack endogenous production are well-known (15). The presence of cheats reduces the fitness of producer cells and puts evolutionary pressure on SP-biosynthesis and receptor genes to reduce such piracy (16). Within the nasal cavity, host lactoferrin restricts the availability of free iron and iron-acquisition genes of *S. aureus* are induced (17). Hence, it seems plausible that SP-production is of importance within this habitat. Additionally, the well investigated routes of iron-acquisition of *S. aureus* suggest adaption towards the presence of siderophores produced by distantly related bacteria (xenosiderophores). *S. aureus* can produce and utilize two different SPs, staphyloferrin A (SF-A) and staphyloferrin B (SF-B) (18,19). In addition, it is able to use hydroxamate and catecholate xenosiderophores such as aerobactin, ferrichrome or bacillibactin (20,21). However, the extent to which *S. aureus* competes for SPs with other members of the nasal microbiome is unknown. Similarly, it is unclear if SP-piracy between *S. aureus* and certain commensals might explain positive or negative correlations between members of the human nasal microbiome.

We have investigated SP-based interactions between *S. aureus* and other members of the nasal microbiome. We tested the ability of 94 nasal isolates from 11 different genera to produce SPs and tested the ability of SP-producers to support *S. aureus* growth. Conversely, we tested the ability of *S. aureus* to support the growth of SP-deficient species in a staphyloferrin-dependent manner. We found a plethora of SP-mediated interactions. Importantly, apart from *C. propinquum,* most *Corynebacterium* species did not produce SPs but were able to consume *S. aureus* SF-A and/or SF-B. In co-cultivation experiments we found that staphyloferrin piracy by competitors created a physiological burden for *S. aureus* and their presence delayed its proliferation in an iron-dependent manner. Finally, we detected siderophores in the nasal cavity of human volunteers and demonstrated that SP-acquisition is crucial for *S. aureus* to proliferate in the cotton rat model of nasal colonization, suggesting that SP-based competition or collaboration might be relevant for structuring the human nasal microbiome.

## Results

### Siderophore production among nasal bacterial species is diverse

It is known that *S. aureus* produces staphyloferrins but can also use siderophores produced by other species (xenosiderophores). We hypothesized that other nasal commensals might possess similar abilities, resulting in reciprocal dependencies.

To investigate this, we tested 94 bacterial isolates derived from the nasal cavity of human individuals in Münster (Germany) (22) as well as in Tübingen (Germany) (23). The collection comprised species from 11 genera namely *Staphylococcus*, *Bacillus*, *Citrobacter*, *Corynebacterium*, *Cutibacterium* (*Propionibacterium), Dolosigranulum*, *Finegoldia, Mammaliicoccus, Moraxella* and *Streptococcus* including several strains from the same species. All strains were screened for siderophore-production using the chrome azurol S overlay assay (O-CAS) (24). The assay indicates SP-production by a color change to yellow as illustrated in Fig. 1A. *Staphylococcus aureus* USA300 LAC generates a yellow color due to the production of staphyloferrins, while *Staphylococcus lugdunensis* N920143 is unable to produce SPs (25). The results for all isolates are summarized in Fig 2. Apart from *S. lugdunensis* and *Staphylococcus hominis,* all staphylococcal strains tested produced SPs. This was expected as the locus *sfaDABC* encoding the genes for SF-A biosynthesis is highly conserved amongst staphylococci (18). In addition, the *Bacillus*, *Citrobacter* and *Mammaliicoccus sciuri* (formerly *Staphylococcus sciuri*) isolates also produced SPs. Interestingly, among the *Corynebacterium* isolates only *C. propinquum* produced siderophores. None of the other isolates including *Dolosigranulum pigrum*, *Finegoldia magna, Moraxella catarrhalis, Peptoniphilus harei,* cutibacteria and streptococci showed positive reactions in the CAS assay.

**Fig. 1:**
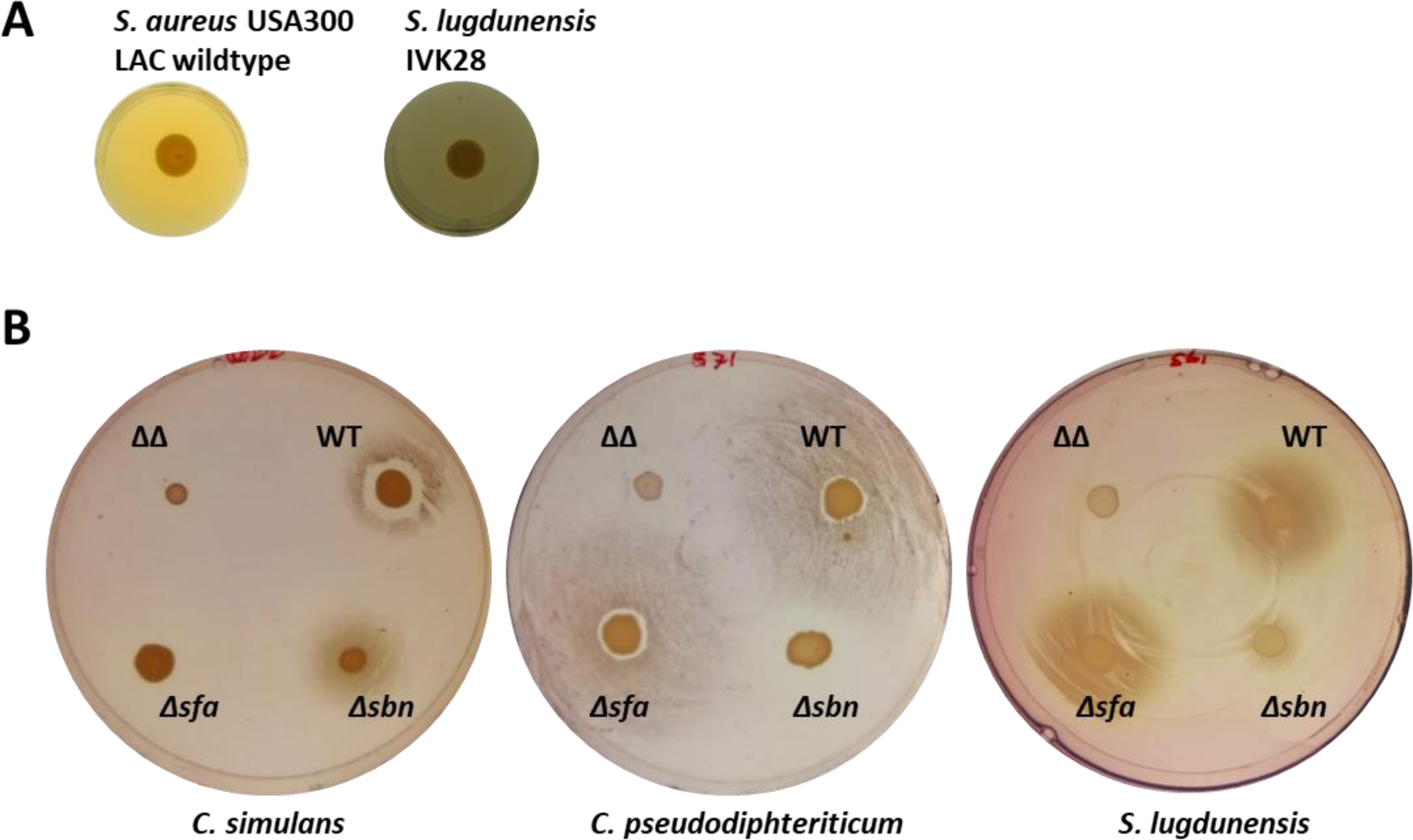
Siderophore production and staphyloferrin usage by nasal bacterial isolates **A) Siderophore production:** Bacterial isolates were spotted on BHI-EDDHA agar in 24 well plates and incubated for one week at 37°C. After incubation wells were overlaid with CAS-containing top agar. Color change to yellow indicates siderophore production and was assessed **4** hours after the overlay. Shown are results for *S. aureus* USA300 LAC and *S. lugdunensis* N920143. |**B) Staphyloferrin A and staphyloferrin B usage by nasal commensals:** An even lawn of nasal commensal species was applied to iron-depleted RPMI plates with 10% horse serum. *S. aureus* strains producing either both staphyloferrins (WT), only staphyloferrin A (Δ*sbn*), only staphyloferrin B (Δ*sfa*) or none of the two (ΔΔ) were spotted on top of the lawn. Growth surrounding the strains was assessed after **24h (*C. simulans*** and ***S. lugdunensis*)** or **48 hours (*C. pseudodiphtheriticum*)** of incubation. Results of *C. simulans* (50Mns_SDm2), *C. pseudodiphtheriticum* (44VAs_Sa4) and *S. lugdunensis* (N920143), are shown. Siderophore-production and staphyloferrin-consumption of all nasal isolates is summarized in Fig. 2.

**Fig. 2:**
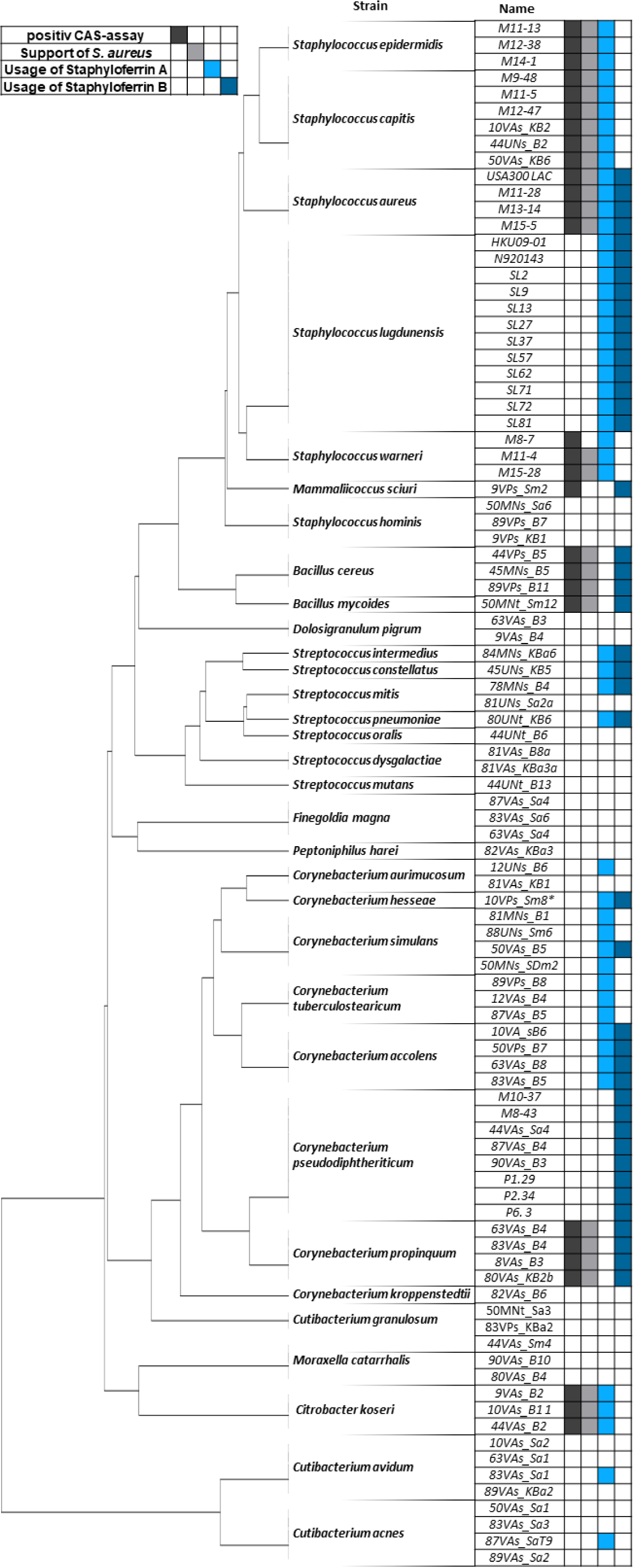
Summary of siderophore production, and consumption. Siderophore production, indicated by positive signals in CAS overlay assays, is shown by dark grey. Growth improvement of *S. aureus* Δ*sbn*Δ*sfa* surrounding siderophore producers is indicated by light grey color. Growth improvement of commensals surrounding *S. aureus JE2* Δ*sbn* and *S. aureus JE2* Δ*sfa* is shown by blue, respectively. * The strain *C. hesseae* 10VPs_Sm8 was initially identified as a *C. aurimucosum* isolate.

### Nasal commensals profit from staphyloferrin producing *S. aureus*

We speculated that nasal organisms lacking endogenous SP production might be able to acquire xenosiderophores. Staphylococci are frequent colonizers of the nasal cavity and prominent producers of siderophores. The majority of human-associated staphylococcal species produce only staphyloferrin A (SF-A) while *S. aureus* produces SF-A as well as staphyloferrin B (SF-B). To test if staphyloferrin-producing bacteria could promote growth of nasal commensals, we constructed isogenic mutants of *S. aureus* JE2 lacking *sfaDABC* (no production of SF-A)*, sbnA-I* (no production of SF-B) or a double mutant (no SP production). Only the double mutant was negative in the CAS assay, confirming complete loss of SP-production (Fig S1). Lawns of nasal commensals were applied to iron-limited agar plates containing 10% horse serum (HS-RPMI). The wild type *S. aureus* and the isogenic mutants were dotted on top of the lawn. After incubation, we often observed growth of the commensals surrounding *S. aureus* WT or individual mutants being stimulated. In contrast, the *S. aureus* double mutant did not stimulate growth of commensals (Fig. 1B and Fig. 2). Growth enhancing effects around *S. aureus* colonies was frequently observed even when commensals (forming the lawn) produced siderophores endogenously. Most staphylococcal species profited from SF-A but not from SF-B production as indicated by improved growth surrounding the *S. aureus Δsbn* but not the *Δsfa* mutant (Fig. 2). *S. hominis* strains were the only staphylococcal species that did not benefit from SF-A or SF-B producing *S. aureus* while *S. lugdunensis* strains were the only s*taphylococci* profiting from SF-A as well as from SF-B producing *S. aureus* (Fig. 1B, Fig. 2). This phenomenon was previously described for *S. lugdunensis* (25). A prominent observation was that the production of SF-A and/or SF-B supported the growth of almost all *Corynebacterium* isolates. With the exception of a single *Corynebacterium aurimucosum* strain and *Corynebacterium kroppenstedtii*, all *Corynebacteria* profited from either SF-A (7 of 27 strains), SF-B (12 of 27 strains) or both (6 of 27 strains). However, *B. cereus*, *B. mycoides*, *C. koseri, M. sciuri*, several s*treptococcal* isolates and individual strains of *Cutibacterium acnes* and *Cutibacterium avidum* were stimulated by staphyloferrin-producing *S. aureus* (Fig. 2).

Altogether, these data indicate that staphyloferrins represent a widely accessible iron source for the bacterial species of the nasal cavity.

### Staphyloferrin consumption by *Corynebacterium hesseae* is receptor dependent

*S. aureus* uses the membrane bound lipoproteins HtsA and SirA to acquire ferric forms of SF-A and SFB, respectively (26–29). We reasoned that nasal commensals might express homologous receptors enabling them to acquire staphyloferrins. To investigate this, we sequenced the genomes of six staphyloferrin-consuming strains from different species. We found genes encoding proteins with homology to HtsA and SirA in all strains (Table S1). However, amino acid sequence identity to the staphylococcal receptors varied between 24% and 62% which seems too low to draw direct conclusions about the substrate specificity. To further investigate this, we focused on the SF-A and SF-B consuming *Corynebacterium hesseae* strain 10VPs_Sm8 which we found amenable to genetic manipulation. The strain encodes the protein R3O64_11615 (43% and 26% identity to SirA and HtsA, respectively) as well as the protein R3O64_03755 (27% and 26% identity to SirA and HtsA, respectively). We created deletion mutants lacking these genes and tested their ability to thrive in the presence of *S. aureus*. Interestingly, we found loss of R3O64_11615 to be sufficient to abrogate growth promotion by SF-A as well as by SFB producing *S. aureus* while loss of R3O64_03755 did not influence this phenotype (Fig.3). This data suggests that SF-A and SF-B usage of *C. hesseae* is mediated by a single receptor. Additionally, these data demonstrate that the growth stimulation was solely caused by staphyloferrins and not by other metabolic products of *S. aureus*.

**Fig. 3:**
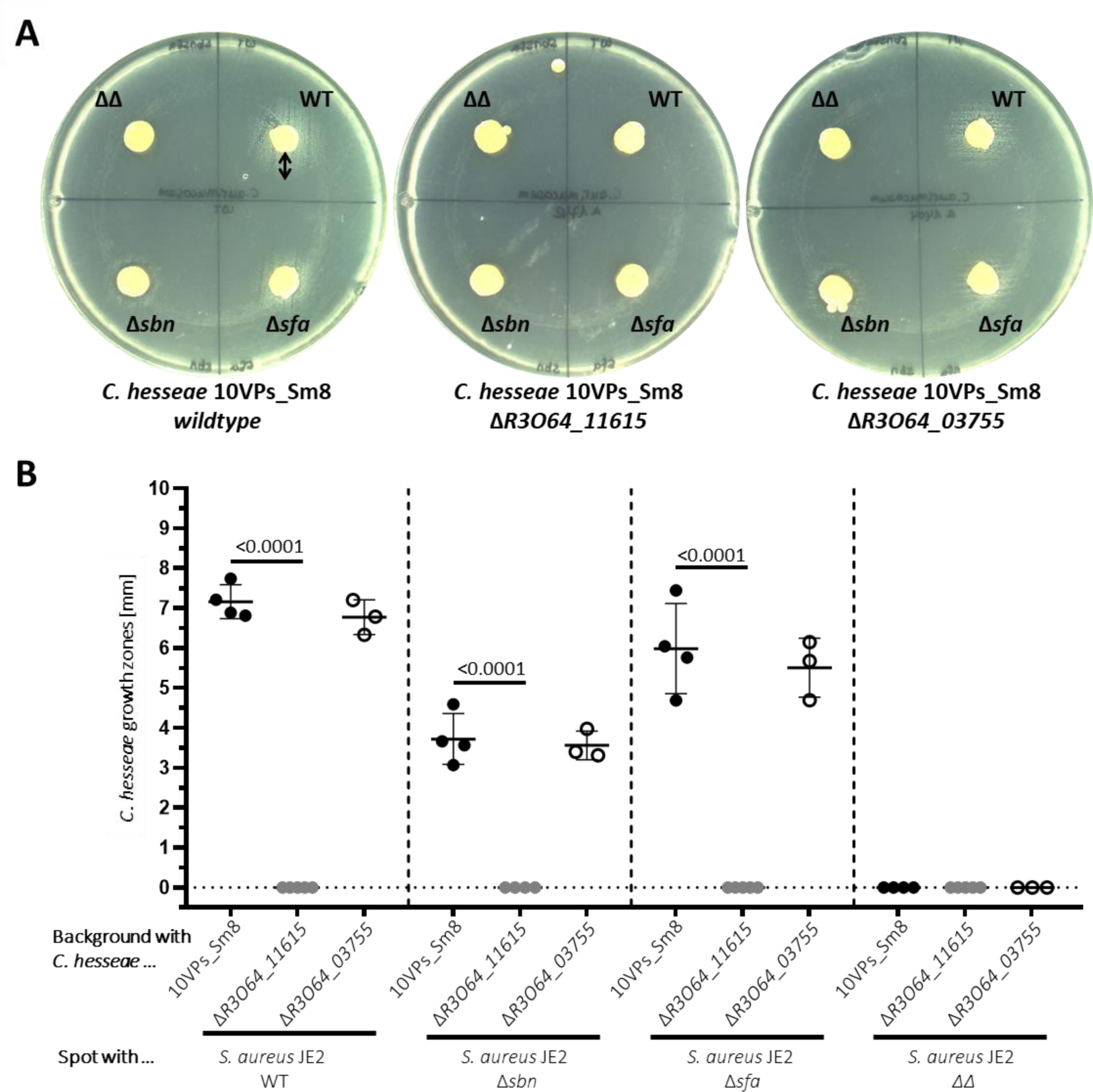
Staphyloferrin consumption by *Corynebacterium hesseae* 10VPs_Sm8. An even lawn of *C. hesseae* (10VPs_Sm8) or its isogenic mutants *R3O64_11615* and *R3O64_03755*) were applied to iron-depleted RPMI plates. *S. aureus* strains producing either both staphyloferrins (WT), only staphyloferrin A (Δ*sbn*), only staphyloferrin B (Δ*sfa*) or none of the two (ΔΔ) were spotted on top of the lawn. Growth surrounding the spots (indicated by black arrow) was quantified 48 h post incubation. **A)** Representative plates of the experiment. **B)** Mean and SD of 3 to 4 independent experiments is shown. Statistical analysis was performed using ordinary one-way ANOVA (<0.0001) with subsequent multiple comparison.

### *S. aureus* profits from siderophore producing commensals

*S. aureus* can acquire xenosiderophores of the hydroxamate-type via the FhuABCD1/D2 system and catecholate type siderophores via SstABC. Therefore, we speculated that *S. aureus* might profit from the presence of SP-producing strains identified.

To test this, we plated even lawns of *S. aureus* Δ*sfa*Δ*sbn* on iron-limited HS-RPMI plates, spotted SP-producing nasal isolates on top of the lawn and assessed *S. aureus* growth surrounding the inocula (Fig. 4A, Fig. 2). Of note, the Δ*sfa*Δ*sbn* mutant does not secrete staphyloferrins but all SP receptors are expressed.

**Fig. 4:**
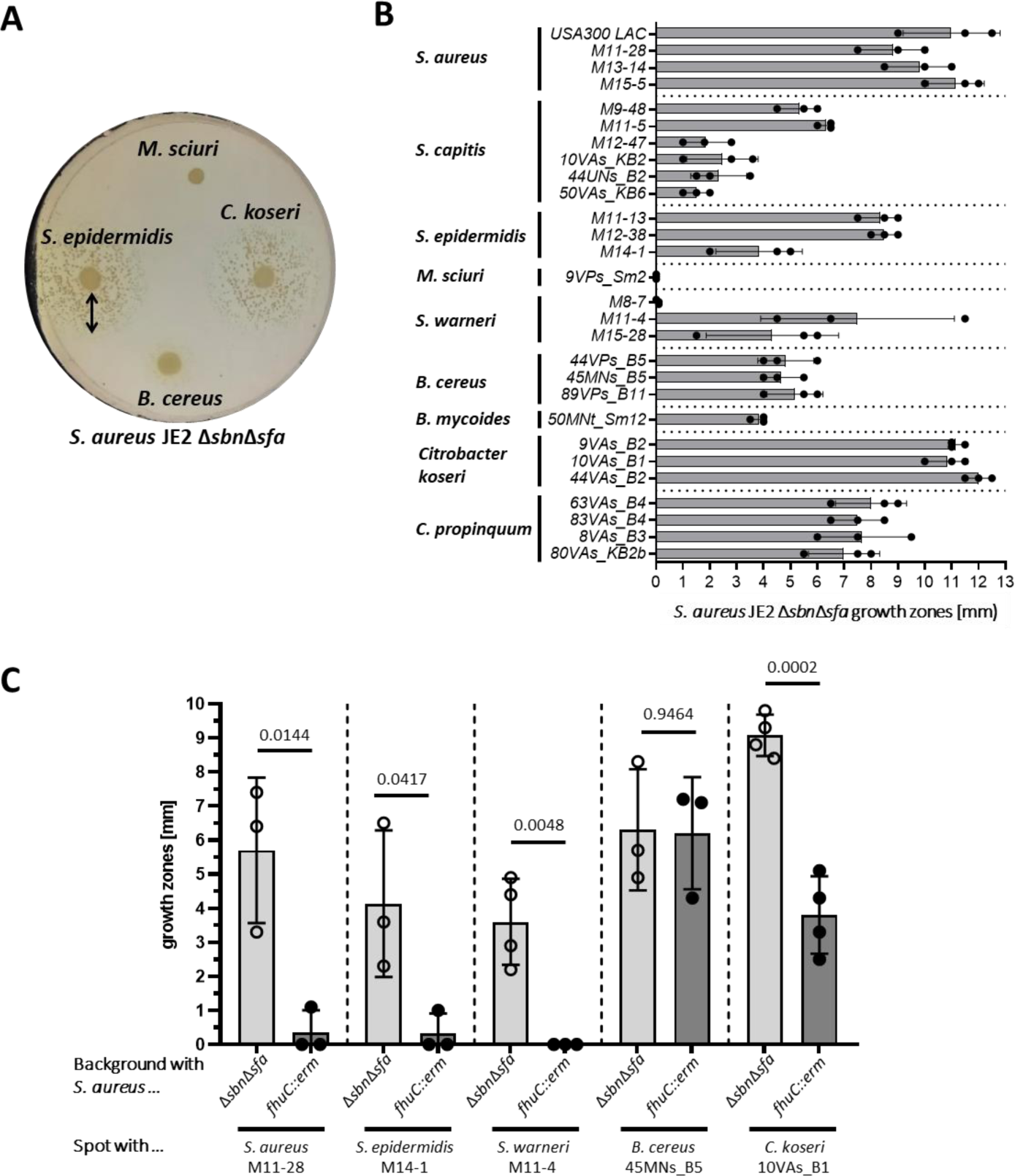
Usage of xenosiderophores by *S. aureus*. An even lawn of *S. aureus* JE2 Δ*sbn*Δ*sfa* or *fhuC::erm* was applied to iron-depleted RPMI plates with 10% horse serum SP-producing nasal isolates were spotted on top of the lawn. Growth surrounding the spots (indicated by black arrow) was quantified 17,5 h post incubation. **A)** An example plate of *S. aureus* JE2 **Δ*sbn*Δ*sfa*** with strains *S. epidermidis* M14-1*; Citrobacter koseri* 10VAs_B1; *B. cereus* 45MNs_B5; *Mammaliicoccus sciuri* 9VPs_Sm8) is shown. **B)** Quantitative evaluation of *S. aureus* Δ*sbn*Δ*sfa* growth zones around different siderophore producers. Mean and SD of 3 independent experiments is shown. **C)** Comparison of growth zones of *S. aureus* JE2 Δ*sbn*Δ*sfa* and *fhuC::erm* surrounding the SP-producers. Mean and SD of 3 to 4 independent experiments is shown. Statistical analysis was performed using unpaired t test.

All *S. aureus* isolates fostered growth of the Δ*sfa*Δ*sbn* mutant which is consistent with the production of SF-A and SF-B. Interestingly, *C. propinquum* and *Citrobacter koseri* isolates promoted proliferation of the Δ*sfa*Δ*sbn* mutant, suggesting that these strains produce siderophores that are accessible to *S. aureus*. Most coagulase negative staphylococcal isolates also allowed growth of *S. aureus* Δ*sfa*Δ*sbn* which is consistent with the previous finding that staphylococci carry SF-A biosynthesis genes (18). Finally, the *Bacillus cereus* and *Bacillus mycoides* isolates allowed proliferation of *S. aureus* Δ*sfa*Δ*sbn*. In contrast, individual isolates of *Staphylococcus warneri and M. sciuri* did not allow proliferation of the *S. aureus* mutant, suggesting that their SPs are not accessible to *S. aureus*. We analyzed the genome sequences of selected SP-producers to identify their siderophores (Tab. S2). *Citrobacter koseri* 44VAs_B2 encoded the biosynthesis genes for aerobactin (mixed type), turnerbactin (catecholate type) and yersiniabactin (phenolate type). Aerobactin has been shown before to support *S. aureus* growth via the FhuABCD1/D2 system (20), and turnerbactin might be acquired via the SstABC system (30). *B. cereus* 45MNs_B5 encoded genes for synthesis of the catecholate type siderophores bacillibactin and petrobactin which might be acquired by *S. aureus* via the SstABC system. In contrast *M. sciuri* 45MNs_B5 which did not support *S. aureus* proliferation encoded a single, yet uncharacterized SPbiosynthesis cluster.

FhuC serves as a house keeping ATPase that energizes acquisition of SF-A, SF-B and hydroxamate siderophores (18,20,31). The catecholate acquisition system SstABC of *S. aureus* is independent of FhuC (30). We used a *fhuC::erm* mutant of *S. aureus* JE2 (32) to verify that the enhanced proliferation of *S. aureus* surrounding the commensal was caused by provision of xenosiderophores. Enhanced growth of the *fhuC* deficient mutant surrounding *S. aureus*, *S. epidermidis* and *S. warneri* strains was not observed (Fig. 4B), further strengthening that staphyloferrins were responsible for growth stimulation. Proliferation of the *fhuC* deficient strain surrounding *Citrobacter koseri* was markedly reduced while growth surrounding *B. cereus* was independent of FhuC Fig. 4C). This data agrees with *Citrobacter koseri* producing both hydroxamate and catecholate type SPs while *B. cereus* produces exclusively cat-echolate type SPs which are acquired independently of FhuC.

These experiments confirm that the growth enhancement is dependent on the receptor-specific exchange of siderophores between *S. aureus* and nasal commensals. Additionally, the data show that increased proliferation of *S. aureus* surrounding nasal commensals depends on its ability to import the produced SPs.

### Classification of siderophore-based interactions between *S. aureus* and nasal commensals

Our experiments show that *S. aureus* interacts with nasal commensals in three distinct fashions (Fig. 5). Firstly, several strains did not produce siderophores but consumed staphyloferrins (cheaters). Most corynebacterial isolates belong to this category (Fig. 5A). Secondly, certain commensal species produced siderophores that are inaccessible to *S. aureus* while simultaneously staphyloferrins are consumed. This interaction is referred to as “locking away” (16) (Fig. 5B). The *M. sciuri* isolate is a prominent example in this type. Thirdly, several commensals produced siderophores that supported *S. aureus* growth while staphyloferrins were consumed by the commensal “shared labor” (Fig. 5C). *Citrobacter koseri* and *Corynebacterium propinquum* belong to this category.

**Fig. 5:**
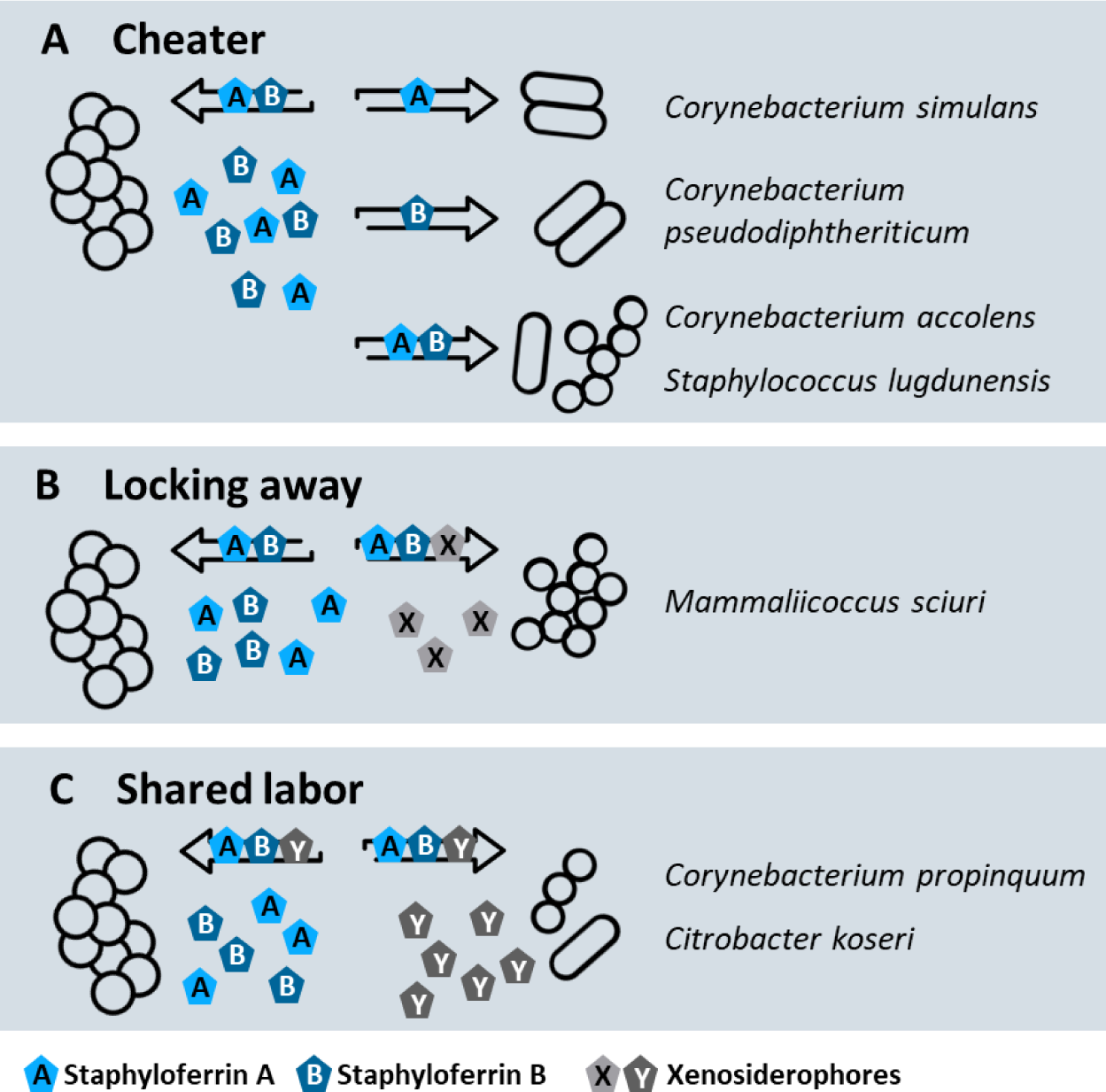
Schematic diagram of siderophore-based interactions between *S. aureus* and nasal commensals. **A)** “Cheater” phenotype. Commensals without endogenous SP-production consume staphyloferrins. **B)** “Locking away” phenotype. Commensals consume staphyloferrins while endogenous siderophores of the commensals are NOT accessible to *S. aureus*. **C)** “Shared labor” phenotype. Siderophores produced by *S. aureus* and the commensals are reciprocally accessible.

Siderophore production is metabolically costly and it is known that SP-based interactions shape bacterial communities (15,16). In particular competitors with the “cheater” and “locking away” phenotype decrease the relative fitness of SP-producers in bacterial communities. This suggests that certain nasal commensals might decrease the fitness of *S. aureus* in iron-limited environments.

### Staphyloferrin-consumption by competitors reduces fitness of *S. aureus*

We designed a physiological assay to investigate if siderophore consumption by competitors might impact the proliferation of *S. aureus* LAC. The staphyloferrin-producing wild type strain was able to proliferate in iron-deficient RPMI with holo-transferrin as a sole source of iron, while the Δ*sfa*Δ*sbn* did not grow (Fig. 6A). Interestingly, the Δ*sfa* mutant showed normal levels of growth while mutation of *sbn* abrogated growth. This indicates that staphyloferrin B is crucial under these experimental conditions (Fig. 6A). This is explained by the fact that exponential growth of *S. aureus* in the presence of glucose is based on fermentation (33,34). Previous studies showed that SF-B production is dominant over SF-A production during fermentation due to a dedicated citrate synthase that allows SF-B production even when activity of the TCA cycle is low (35,36).

**Fig. 6:**
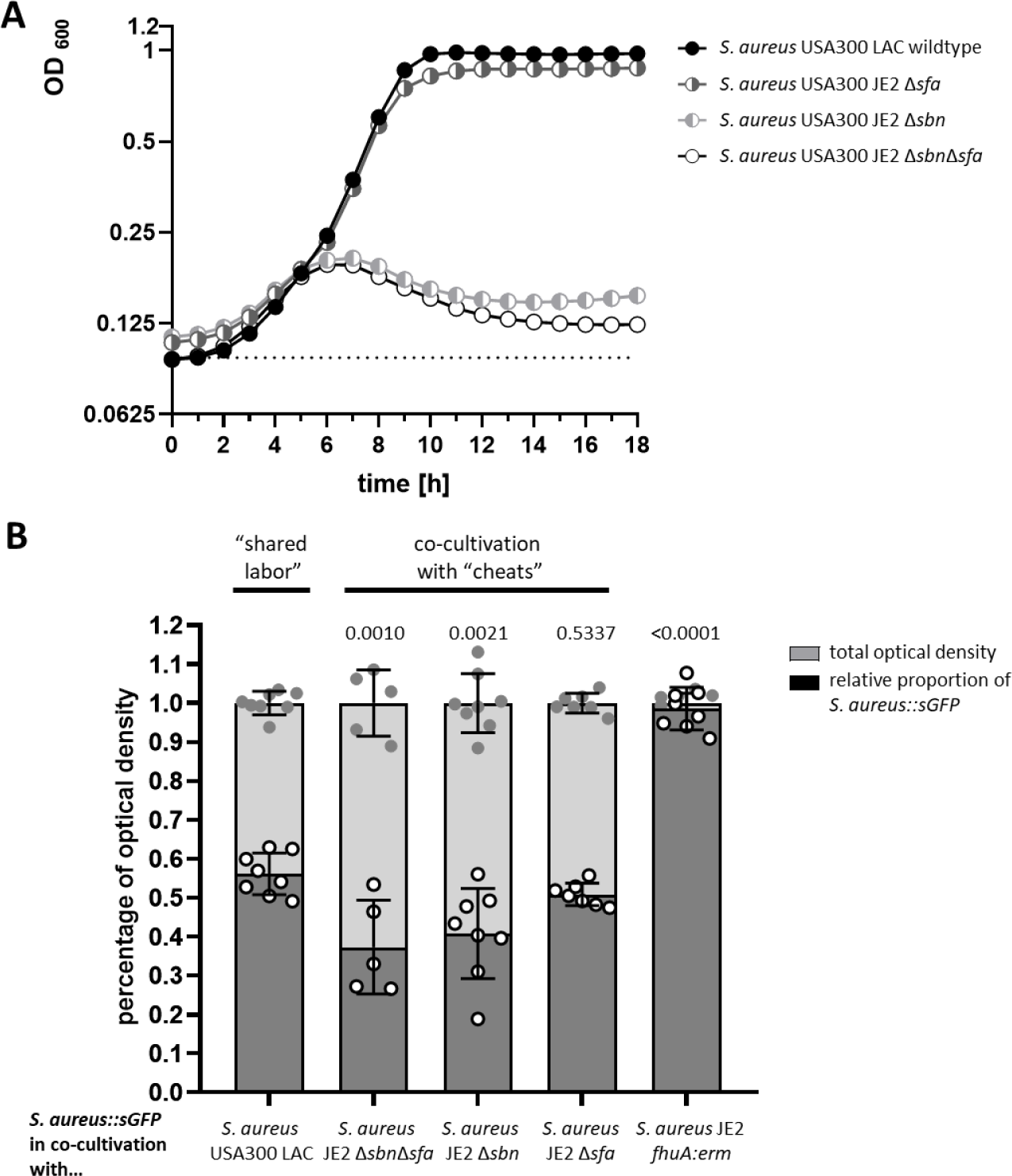
Growth and co-cultivation in iron-limited medium. **A) Growth curves in iron-limited medium:** *S. aureus* USA300 LAC wildtype and *S. aureus* strains lacking either the staphyloferrin A and B or both production genes were inoculated to an optical density OD 0.01 and grown for 18 h in 500 µl iron-limited medium (1x RPMI, 1% casamino acid, 10 µM EDDHA, 100 µg holo-transferrin) at 37°C under constant shaking in the Tecan Spark ® 10M multimode microplate reader. Growth was monitored via optical density at 600 nm. Shown is the mean of three to ten biological replicate of each strain. The dotted line indicates the media control. **B) Co-cultivation in iron-limited medium:** For each co-cultivation the strain *S. aureus::sGFP* was mixed in a 1:1 ratio (start OD of 0.02) with *S. aureus* USA300 LAC wildtype, *S. aureus* lacking staphyloferrin production genes and *S. aureus* defective in its siderophore uptake systems (*fhuC::erm*). The strains were grown for 24 h in 500 µl iron-limited medium at 37°C under constant shaking in the Tecan Spark ® 10M multimode microplate reader. Growth was monitored via optical density at 600 nm and fluorescence intensity of the GFP signal (Ext. 480 nm and Em. 525 nm). The fluorescence values were extrapolated to the optical density values by using the formula y=12.54+1284*x. The extrapolated values are shown in dependence of the measured optical density values at timepoint of 10 h, which were set to 1. Statistical analysis was performed using ordinary one-way ANOVA (<0,0001) with subsequent multiple comparison by comparing all relative proportions of *S.aureus::sGFP* of each mixed culture with the shared labor condition.

For competition experiments we used an *S. aureus* LAC expressing sGFP (37).This did not influence the growth rate of the strain (Fig. S2A) but produced a clear fluorescent signal in linear proportion to the optical density of the culture (Fig S2B, C).

We mixed the wild type *S. aureus* expressing sGFP with unlabeled *S. aureus* wild type or staphyloferrindeficient mutants in an equal ratio to reflect conditions of “shared labor” and “staphyloferrin cheating”, respectively. Analysis of the GFP/OD correlation was used to assess the proportion of both strains at the end of the experiment. “Shared labor” resulted in even ratios of GFP positive to GFP negative cells (Fig. 6B), indicating that both strains grew equally well. In contrast, competition with the Δ*sbn*Δ*sfa* and Δ*sbn* mutants allowed similar OD values to be reached but reduced the GFP signal of the mixed culture, suggesting decreased fitness of the WT compared to the cheats. Competition with the SF-A deficient strain did not affect fitness of the WT which correlates with the observation that SF-A production was dispensable under these conditions. To further confirm the relevance of siderophoresharing, we included the *fhuC* deficient strain (USA300 JE2 *fhuC::erm*). The strain is able to produce SFA and SF-B but fails to acquire them. The *fhuC::erm* mutant was completely displaced by the GFPpositive WT strain by the end of the experiment, highlighting the relevance of SP-acquisition in the competition experiment. Together, these experiments show that siderophore usurpation by competitors reduces the competitive fitness of staphyloferrin producing *S. aureus* strains.

In this light, we hypothesized that the presence of nasal isolates that consume staphyloferrins or provide xenosiderophores might impact the growth of *S. aureus* during co-cultivation. Nasal commensals failed to grow under the liquid co-culture conditions described above. Therefore, we used agar plate-based experimental conditions and chose individual nasal strains showing different interactions with *S. aureus*. Firstly, we chose the “cheater” *C. pseudodiphtheriticum* which consumes SF-B and investigated its effects on *S. aureus* USA300 JE2. Mono as well as mixed cultures (1:1 ratio) were plated on iron limited HS-RPMI and the size of *S. aureus* colonies was measured after 24 h of incubation (Fig.7). *S. aureus* formed regular sized colonies in monoculture (mean colony size of 0,4188 mm) while *C. pseudodiphtheriticum* was unable to form colonies under these conditions. During co-culture the size of the *S. aureus* colonies was reduced by 60% (mean 0.168 mm) although *C. pseudodiphtheriticum* colonies were hardly detectable. To verify that this effect was caused by competition for iron, we included 20 µM FeSO_4_ into the plates which increased the mean colony size of *S. aureus* in co-culture to 1,045 mm. This represents an increase of 160% above the level achieved in monoculture in the absence of additional iron. The additional iron also allowed *C. pseudodiphtheriticum* to form visible colonies. This strongly suggests that iron-limitation is the dominant driver of *S. aureus* colony size in this assay.

**Fig. 7:**
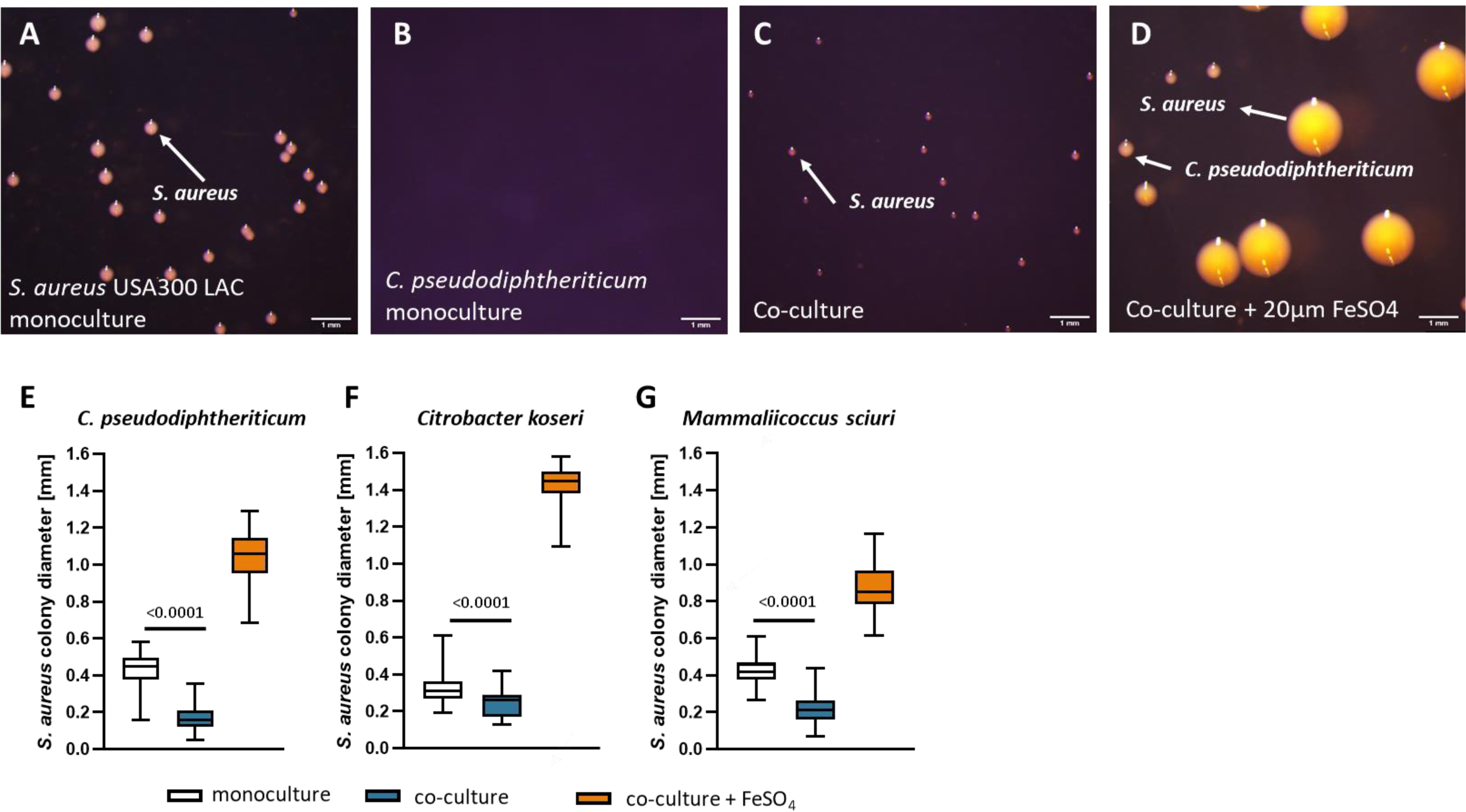
Siderophore-based interactions with nasal commensals impact growth of *S. aureus*. **A-E)** *S. aureus* USA 300 LAC wildtype and *C. pseudodiphtheriticum* strain 90VAs_B3 were either plated individually (A+B), or mixed in even numbers (C-D) and plated on **HS-RPMI** plates (A-C) or **HS-RPMI** plates **supplemented** with **20 µM FeSO_4_** (D) After **24 h** incubation at 37 °C, pictures of the plates were taken and the diameter of *S. aureus* colonies was measured using ImageJ. Representative pictures are shown. **E-G)** Summary of *S. aureus* colony sizes under the various conditions and cocultures with *C. pseudodiphtheriticum* strain 90VAs_B3, *Citrobacter koseri* 44VAs_B2 and *Mammaliicoccus sciuri* 9VPs_Sm2. Data represent the diameter of 44-232 individual colonies from three independent experiments. Floating bars represent 25 and 75% percentiles, the horizontal lines represent the medians. Whiskers show minimum and maximum range. Statistical analysis was performed using unpaired Mann-Whitney t-test.

We speculated that species “sharing labor” with *S. aureus* by providing accessible siderophores might not impact *S. aureus* colony formation. To test this, we used *C. koseri* (strong support of *S. aureus* – Fig. 4) and repeated the co-culture experiment. We found that that the presence of. *C. koseri* reduced *S. aureus* colony size only slightly (16%) suggesting less fierce competition between the species. *M. sciuri* produces a siderophore which is not accessible to *S. aureus* while simultaneously consuming SF-B (locking away interaction). Co-culture reduced the size of *S. aureus* colonies by 49%. This is indication of strong competition but the effect was not more severe than that observed for co-culture with *C. pseudodiphtheriticum*.

### Siderophore acquisition is needed for nasal colonization by *S. aureus*

To investigate the relevance of siderophore acquisition to the ability of *S. aureus* to colonize the nares of cotton rats, we replaced the wild type *fhuC* gene of a streptomycin resistant mutant of *S. aureus* Newman with a *fhuC::erm* mutation (32). As described previously, the Δ*fhuC* mutant grew normally in the presence of FeSO_4_ but was strongly disabled in iron-limited medium even if aerobactin (a substrate of FhuBGD_1_) was added. Similarly, the mutant grew poorly when culture supernatants of *S. aureus* Δ*sfa* or Δ*sbn* were added as sources of SF-B or SF-A, respectively which has been described before (38). All phenotypes could be complemented by plasmid-based expression of FhuC (Fig. S3).

We used the cotton rat model of nasal colonization to investigate the relevance of SP-acquisition for nasal colonization. Compared to the WT, the level of colonization was significantly reduced when the *fhuC::erm* mutant was implanted (Fig 8A). This confirmed the iron-limited environment within the nasal cavity and shows that SP-acquisition is relevant for *S. aureus* to overcome the nutritional iron-limitation within the nasal cavity.

**Fig. 8:**
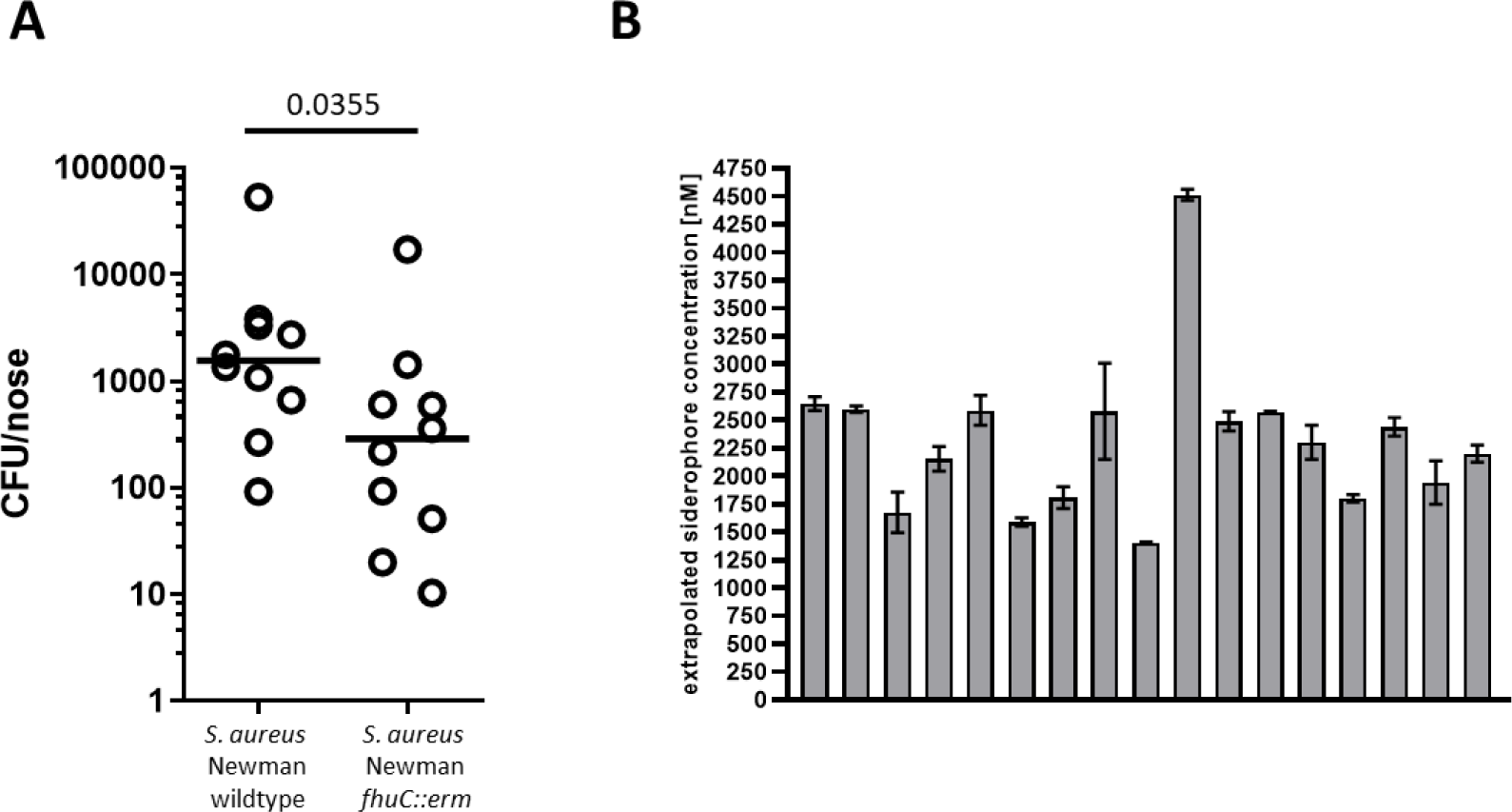
Siderophore acquisition important for nasal colonization of cotton rats and iron chelating activity is identified in human nasal specimen. **A) Cotton rat nasal colonization.** 1x10^7^ CFU of *S. aureus* strain Newman^strepR^ and the isogenic *fhuC::erm* mutant were inoculated into the nares of 8-12 week old cotton rats. 5 days post inoculation the animals were sacrificed, noses were harvested and *S. aureus* CFU were enumerated. Statistical analysis was performed using unpaired Mann-Whitney t-test. **B) Siderophore level in the human nose.** Nasal specimens of 17 random volunteers were collected and the iron-chelating activity was determined using the SideroTec-HiSens^TM^ Assay (Accuplex Diagnostics Ltd.). Shown are the mean and SD of two technical replicates for each volunteer.

### Siderophores are produced in the nasal cavity of humans

We sampled the nasal cavities of 17 human volunteers and used the Sidero-Tec -HiSens Assay to assess the iron-chelating activity. The measured iron chelating capacity suggested siderophore concentrations of 1.2 – 4.4 µM (Fig. 8B), highlighting that production and most likely competition for siderophores is relevant within the human nasal microbiome.

## Discussion

The production of iron-binding siderophores (SPs) facilitates the acquisition of nutritional iron and is of enormous biological importance for bacteria. SP-production occurs with bacteria that have been isolated from many different environmental habitats including sea and fresh water (39,40), soil (24,41), and also from organisms living in association with multicellular organism including plants and animals (42,43).

Siderophores are important facilitators of microbial interactions. They are secondary metabolites that are resource-intensive to produce. Upon secretion, SPs represent public goods that are accessible to the entire microbial community and many bacterial species possess receptors for such xenosiderophores (20,25,44,45). Several mechanism of positive and negative interference have been described (16). Acquisition of xenosiderophores is usually of benefit to the consumer as iron needs are satisfied without costly production of SPs (25,46). However, sole dependence on xenosiderophores harbors the risk of high dependency on other bacteria. In fact, it has been demonstrated that growth of otherwise “unculturable bacteria” can be promoted by SPs of co-occurring species, exemplifying the existence of organisms that are fully dependent on SPs produced by others (47). Regarding the producer organism, consumption of siderophores by co-occurring non-producers (cheaters) reduces fitness, as the benefits of the costly production are shared with others (46,48). In contrast, some bacteria can reciprocally exchange siderophores (shared labor) which will not skew competitive fitness. Finally, production of uncommon SPs can be advantageous for the producer, since no putative competitor can benefit from its production. In this case the siderophore will cause increased iron restriction for the non-producer (locking away) (49,50). Accordingly, SP-based interactions can foster competition as well as collaboration and influence the structure of bacterial communities. These ecological concepts are frequently studied in environmental communities (15). However, it is becoming more evident that they also apply to microbiomes and can be associated with the susceptibility of the host towards pathogen colonization and infection, a concept applying to plant as well as animal pathogens (43,45,51).

In humans, the immune system potentiates natural iron restriction by production of iron chelating molecules, a phenomenon called nutritional immunity. Accordingly, siderophores are virulence factors enabling the proliferation of pathogens in normally sterile tissue (52). However, nutritional immunity is also acting on mucosal and skin surfaces. Iron-binding lactoferrin is found in human secretions including sweat, tears, breastmilk as well as gastric, pancreatic and nasal secretions (53), strongly suggesting that microbial communities colonizing human body surfaces face severe iron limitation. In fact, characterization of the human gut microbiome showed that changes in nutritional iron intake impose drastic effects on its composition, strongly suggesting that iron availability determines the success of species in polymicrobial communities (42). This is supported by the finding that the probiotic *Escherichia coli* Nissle strain displaces the pathogenic *Salmonella enterica serovar Typhimurium* in a sidero-phore-dependent manner (51). Similarly, xenosiderophore usage is crucial for *Bacteroides thetaiotamicron* to grow in the context of the inflamed gut (45).

Little is known regarding the importance of iron for the human nasal microbiome. However, gene expression analysis of *S. aureus* isolated directly from human as well as from cotton rat nasal cavities has shown that iron acquisition genes are expressed during colonization, suggesting that the bacteria experience iron restriction (17,54,55). In line with this, we showed here that iron-chelating activity is detected within the nasal cavity of humans and that a *fhuC* deficient *S. aureus* mutant was attenuated during nasal colonization of cotton rats. This shows the importance of SP production and acquisition in the nasal cavity. Of note, a *fhuC-* deficient strain is still able to acquire catecholate-type xenosiderophores highlighting that reduction of the acquirable spectrum of SPs reduces the fitness of *S. aureus*.

A similar siderophore-dependency during nasal colonization of mice has been observed for *Klebsiella pneumonia* (56), suggesting a broad relevance of nutritional immunity for reducing pathogen colonization.

Little is known about the network of SP-based interactions within the nasal microbiome. We found that about one third of nasal isolates produce siderophores with dominant producers being staphylococci along with *Corynebacterium propinquum*, *Citrobacter koseri* and *Bacillus cereus*. However, *Citrobacter* and *Bacillus* are not regarded as frequent nasal commensals. Most likely these species are intermittently introduced to and lost from the nasal microbiome, and their role in shaping communities by providing siderophores is therefore unclear. A striking finding of our experiments is that corynebacterial species were prominent consumers of SF-A or SF-B or both. According to the ecological considerations detailed above, this “cheating” phenotype should be associated with negative effects for staphyloferrin producers, which we confirmed *in vitro* and it can be assumed that these interactions are relevant *in vivo*. Studies investigating the composition of the human nasal microbiome have frequently reported connections between *S. aureus* and corynebacterial species. Importantly, the presence of *Corynebacterium* spp. is associated with decreased absolute numbers of colonizing *S. aureus* (5,57,58). This might in part be due to siderophore cheating. Wos-Oxley et al. reported a correlation between *S. aureus* and *C. pseudodiphtheriticum* (6). We found *C. pseudodiphtheriticum* to profit from SF-B which might explain its frequent association with *S. aureus* in human nasal communities. Interestingly, human trials showed that instillation of corynebacterial species (among them *C. pseudodiphtheriticum)* into the nostrils of human volunteers reduced or even abolished *S. aureus* colonization (59,60), suggesting that the species are able to interfere with *S. aureus* colonization *in vivo*. However, to determine if interference is caused by siderophore cheating awaits experimental validation. In our experiments, the only corynebacterial species producing a siderophore was *Corynebacterium propinquum*. This SP was previously described as dehydroxynocardamine and was found to inhibit the growth of *S. epidermidis*, most likely by reducing the availability of nutritional iron. In contrast, *S. aureus* was not affected by dehydroxynocardamine (50). Interestingly, we found that *S. aureus* profited from *C. propinquum*, while *C. propinquum* profited from SF-B but not from SF-A production. This suggest a reciprocal adaption between *S. aureus* and *C. propinquum* resulting in “shared labor” phenotypes. In contrast *S. epidermidis* and *C. propinquum* engage in “locking away” type of interaction as the siderophores are reciprocally inaccessible.

In the experiments reported here we identified multiple strains of different species and genera that interacted with *S. aureus* in diverse fashions (shared labor / cheat / locking away). Interestingly, we found that co-culture under iron limited conditions always resulted in *S. aureus* having a decreased colony size. Most likely this reflects the competition for essential nutrients including carbon and energy sources which are all limited in the experimental conditions employed. However, the “cheating” and “locking” away phenotypes had a more pronounced impact on *S. aureus* colony sizes than “shared labor” which is in line with general ecological principles (16). In a co-submitted manuscript by Rosenstein et al. we demonstrate siderophore-based interactions between *S. epidermidis* and *S. lugdunensis* that can, at least partly, explain the positive correlation of the species within the nasal microbiome of humans.

Of note, in many instances SP-consumption and production phenotypes were strain-rather than species-dependent. Similarly, it has been reported that corynebacteria inhibit *S. aureus* strains with different *agr* alleles to variable degrees (57). This phenotypic heterogeneity suggests that any antagonizing/stimulating effects might be dependent on the genotypes of particular strains and extrapolation of findings to the species level might not always be valid. This has also to be considered for other traits that influence bacterial interactions such as bacteriocin production which are frequently strain dependent (7). This study shows a high diversity of siderophore-based interactions. Staphyloferrins were shown to be important SPs that are consumed by strains of different species and genera. This suggest that commensals can create a hostile environment for staphylococci by siderophore cheating. However, it remains unclear to what extent SP-based interactions influence the composition of the nasal community *in vivo*. We did not perform co-colonization experiments with cotton rats as this model relies on animals harboring a rodent nasal microbiome which is not comparable to humans (61). Animals raised in our facility carry a range of staphylococci, and corynebacterial species along with *Escherichia coli* and *Enterobacter* species (data no shown). Many of these strains produce siderophores (data not shown) that would distort the experiments and prevent meaningful interpretation. In the future studies, to investigate the interactions within the nasal microbiome, the development of a colonization model in gnotobiotic (germ-free) animals is needed to allow the creation of humanized nasal microbiomes *in vivo*.

## Conclusions

We have demonstrated a multitude of diverse siderophore-dependent interactions among members of the nasal microbiome. Several corynebacterial species consume staphyloferrin B which is exclusively produced by the pathogen *S. aureus* showing a particular adaption of the former to the presence of the potential pathogen. Siderophore piracy reduces fitness of the producer. Accordingly, our data provide a mechanistic explanation of the frequently observed reduced level of *S. aureus* colonization in individuals carrying high numbers of corynebacteria and pave the way for the development of nasal probiotics to reduce *S. aureus* colonization in humans.

## Methods

### Chemicals

If not stated otherwise, reagents were purchased from Sigma.

### Bacterial strains, media, and culture conditions

A list of plasmid and bacteria strains used/generated are provided in Table 1 and S3. Additionally, Table S3 provides information about incubation time and media required for the different strains. In general strains were streaked out on blood plates, either BM or TSA and stored at 4°C. Antibiotics were added where appropriate: kanamycin (50 µg/ml), chloramphenicol (10 µg/ml), streptomycin (250 μg/ml).

**Table 1.** Bacterial strains and plasmids.

| Strain/ Plasmid | Genotype/ Description | Source |
| --- | --- | --- |
| **Bacterial strains** |  |  |
| *S. aureus* USA300 LAC | wild type | (62) |
| *S. aureus* USA300 JE2 | Plasmid-cured strain LAC | (32) |
| *S. lugdunensis* HKU09-01 | wildtype | (63) |
| *S. lugdunensis* N920143 | wildtype | (64) |
| *S. aureus* JE2 *fhuC::erm* | strain NE406_SAUSA300_0633 of the Nebraska transposon mutant library | (32) |
| *S. aureus* Newman<sup>strepR</sup> | Streptomycin resistant wild type strain | (8) |
| *S. aureus* Newman<sup>strepR</sup> *fhuC::erm* | Contains the *fhuC::erm* mutation from NE406_SAUSA300_0633 | This study |
| *S. aureus* Newman<sup>strepR</sup> *fhuC::erm* pRB473:*fhuC* | Plasmid based complementation of *fhuC* | This study |
| *S. aureus* USA300 JE2 Δ*sbn*Δ*sfa* | Markerless deletion of Δ*sfaDABC* and Δ*sbnABCDEFGHI* | This study |
| *S. aureus* USA300 JE2 Δ*sfa* | Markerless deletion of Δ*sfaDABC* | This study |
| *S. aureus* USA300 JE2 Δ*sbn* | Markerless deletion of Δ*sbnABCDEFGHI* | This study |
| *S. aureus* USA300 LAC::*sGFP* | sGFP fluorescence marker inserted between the genes *NWNM29-30* | (37) |
| *Corynebacterium hesseae* 10VPs_Sm8 | Isolate from the nasal cavity of a human individual in Münster (Germany) | (22) |
| *C. hesseae* 10VPs_Sm8 ΔR3064_11615 | Markerless deletion of R3064_11615 | This study |
| *C. hesseae* 10VPs_Sm8 ΔR3064_03755 | Markerless deletion of R3064_03755 | This study |

| Plasmids |  |  |
| --- | --- | --- |
| pIMAY | Thermosensitive vector for allelic exchange | (65) |
| pRB473:*fhuC* | Expression of *fhuC* by its native promotor | This study |
| pJSC232 | Kanamycin resistance/sucrose sensitivity vector for allelic exchange in *Corynebacteria* spp. | (66) |

### Transduction of the *fhuC::erm* mutation

*S. aureus* Newman^strpR^_*fhuC::erm* was created using phage transduction (phage Φ11) from the Nebraska transposon mutant library (strain NE406_SAUSA300_0633) into the *S. aureus* Newman^strpR^ background using standard protocols.

### Growth curve analysis of *S. aureus* Newman^strpR^ Δ*fhuC::erm*

Staphylococcal deletion mutant strains were grown overnight in TSB at 37°C with agitation. Cells were harvested and washed with RPMI containing 1% casamino acids (CA) (Difco) and 10 μM ethylenediamine-di(o-hydroxyphenylacetic acid) (EDDHA; fluorochem). OD_600_ was adjusted to 1 and 2.5 μl were mixed with 500 μl of RPMI + 1% CA + 10 μM EDDHA per well (final OD_600_ of 0.005) in a 48 well microtiter plate (Nunc, Thermo Scientific). 200 ng/ml aerobactin (EMC Microcollections GmbH), 200 ng/ml ferrichrome (EMC Microcollections GmbH), 5.7% spent medium from USA300 JE2_ Δ*sbn* containing SF-A, 9,1% spent medium from USA300 JE2_Δ*sfa* containing SF-B, or 20 μM FeSO4 were added as iron sources. OD_600_ was measured every 15 min for 24 h in an Epoch2 reader (BioTek) at 37°C orbital shaking.

### Construction of *sfaDABC* and *sbnH-I* deficient mutants

In-frame deletion in *S. aureus* were generated based on the technique described by Monk et al. (65). The 500 bp sequences up- and downstream of the deletion genes were amplified using primers A and B (A/B) and primers C/D respectively (Table 2). Then the PCR fragments were fused by overlap extension PCR and cloned into pIMAY by restriction digestion and transformed into *E. coli* SA08B. Plasmids were confirmed by DNA sequencing. The plasmids were then electroporated into USA300 JE2 and allelic replacement was performed using standard procedure (29). Mutants were validated by PCR amplification and sanger sequencing of the region of interest.

**Table 2.**
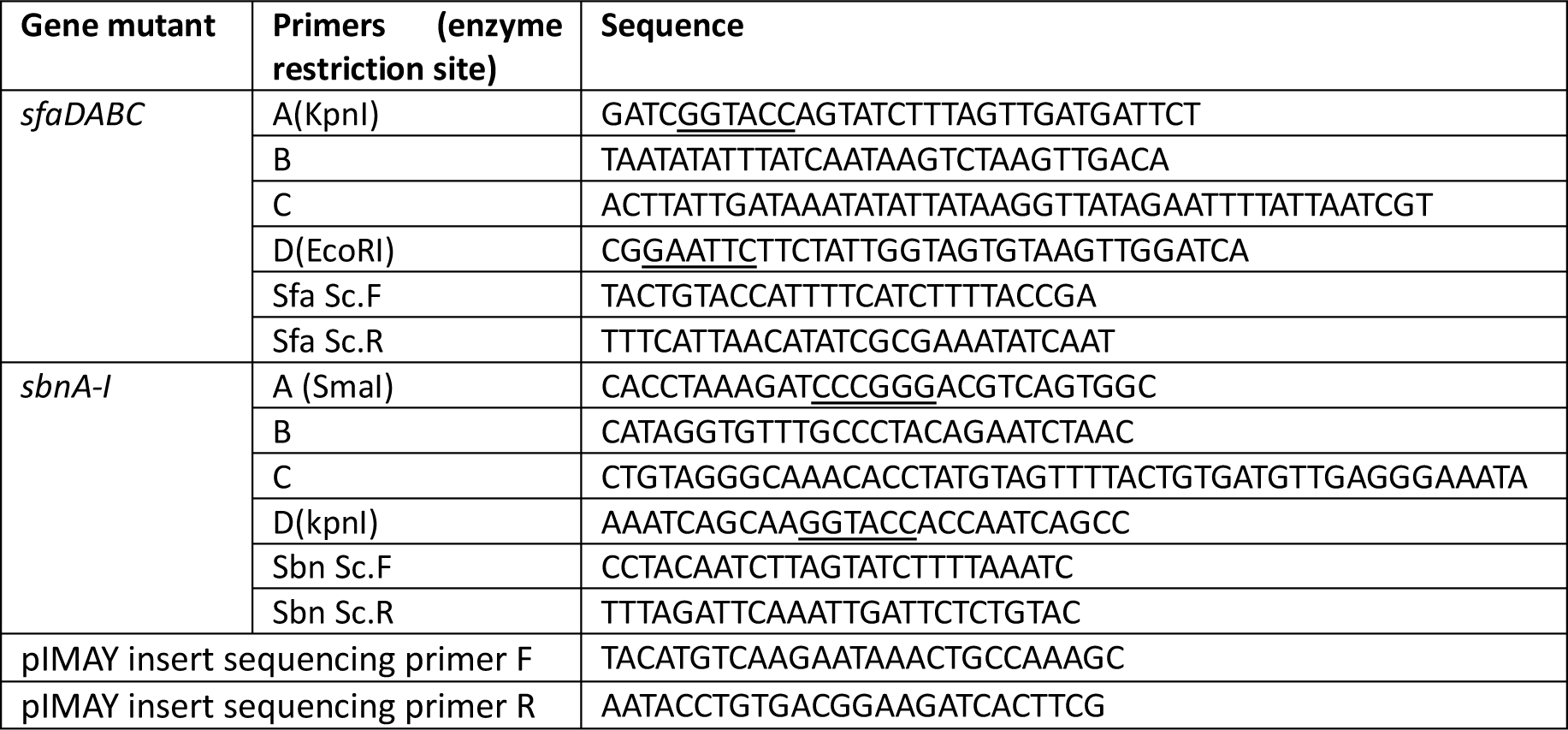
Primers for Construction of *sfa* and *sbn d*eficient Mutants.

### Growth curve analysis of *S. aureus* USA300 JE2 Δ*sfa*, *S. aureus* USA300 JE2 Δ*sbn S. aureus* USA300 JE2 Δ*sbnsfa*

Staphylococcal deletion mutant strains and wild type were grown overnight in TSB at 37°C with agitation. Cells were harvested and washed with RPMI containing 1% CA and 10 μM EDDHA. OD_600_ was adjusted to 1 and 5 μl were mixed with 500 μl of RPMI + 1% CA + 10 μM EDDHA+100 µg holo-transferrin per well (final OD_600_ of 0.01) in a 48 well microtiter plate (Nunc, Thermo Scientific). Holo-transferrin serves as iron source, which allows to see the siderophore dependent growth. OD_600_ was measured every 30 min for 24 h in a Tecan Spark microplate reader at 37°C orbital and linear shaking.

Construction of *R3O64_11615* and *R3O64_03755* deficient *C. hesseae* mutants.

We constructed an in-frame deletion in *C. hesseae* as described previously (66). Upstream and downstream sequences of the target genes were amplified with primers A and B (A/B) and primers C/D, separately (Table 3). The PCR products were fused into the PstI-opened pJSC232 plasmid by sequence and ligation independent cloning (SLIC) and then transferred into *E. coli* DH5α. The insertion was confirmed by DNA sequencing. The plasmids were then electroporated into *C. hesseae* and allelic replacement was performed using kanamycin resistance marker and a negative selection marker for sucrose sensitivity (66). Mutants were validated by PCR amplification and sanger sequencing of the region of interest.

**Table 3.**
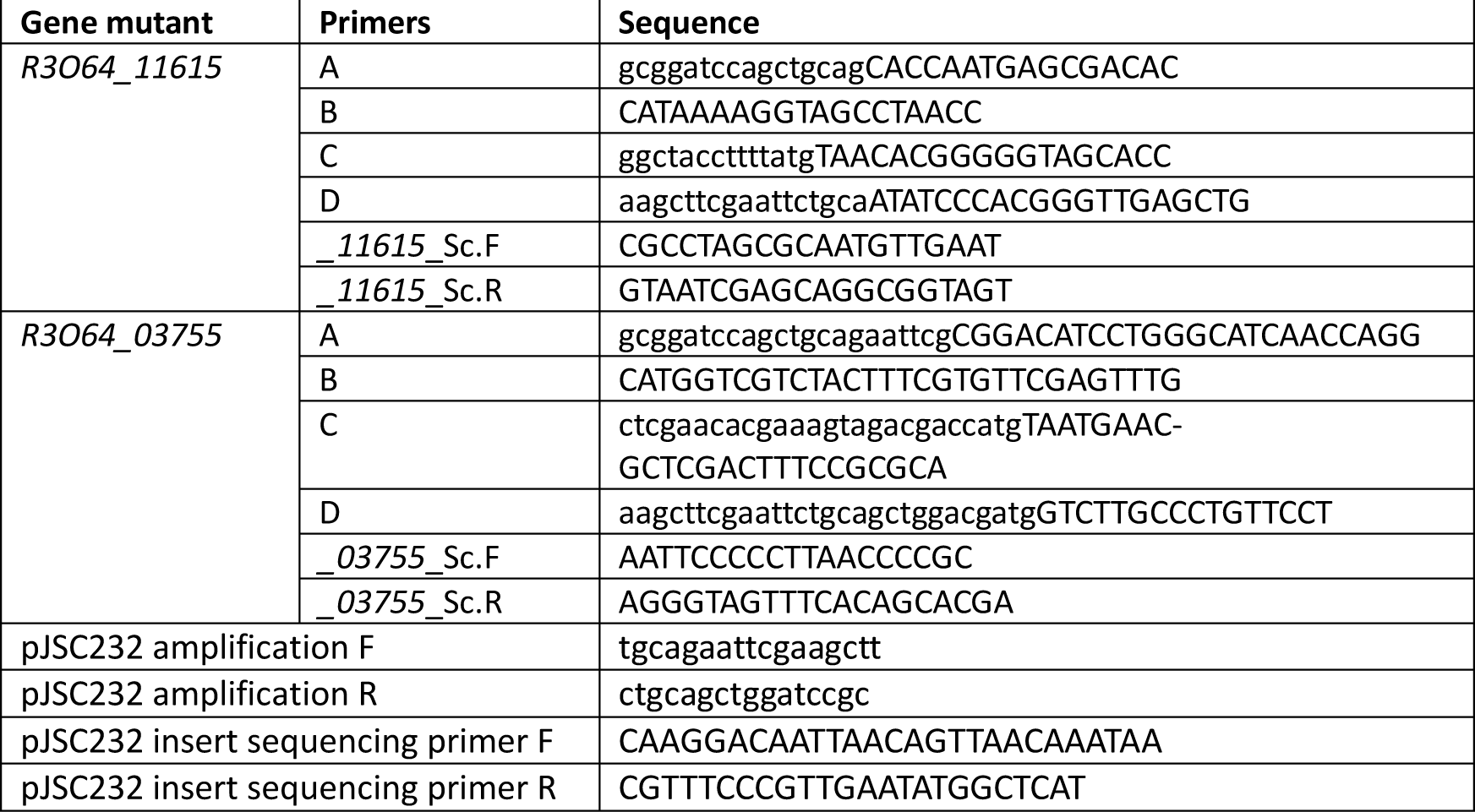
Primers for construction of R3O64_11615 and R3O64_03755 deficient C. hesseae mutants.

### Chrome azurol S overlay assay

The assay was performed as described earlier, with minor modifications (24). For the assay test organisms were grown for one week on BHI-Agar plates containing 10 µM EDDHA and afterwards overlaid with CAS medium. CAS medium was prepared from four solutions which were sterilized separately. Solution 1 (S1) was prepared by mixing 10 ml of 1 mM FeCl_3_*6H_2_0 [in 10mM HCl) with 50 ml of an aqueous solution of Chrome azurol S (CAS) (1.21 mg /ml) and 40 ml of an aqueous solution of hexadecyltrimetyl ammonium bromide (HDTMA) (1.82 mg/ml). Solution 2 (S2) contained Piperazine-1,4bis(2-ethanesulfonic acid) (PIPES) 30.24 g/L in 750ml of a salt solution containing 0.3 g KH_2_PO_4_, 0.5 g NaCI, and 1.0 g NH_4_Cl. The pH was adjusted using 50% KOH to 6.8, then 15 g/l agar was added and the volume was filled up to 800 ml. Solution 3 consists 2 g glucose, 2 g mannitol, 493 mg MgSO_4_*7H_2_0, 11 mg CaCI_2_, 1.17 mg MnSO_4_*H20, 1.4 mg H_3_BO_3_, 0.04 mg CuSO_4_* 5H_2_0, 1.2 mg ZnSO_4_*7H_2_0, and 1.0 mg

Na_2_MoO_4_*2H_2_0 in 70 ml water. Solution 1-3 were autoclaved and then cooled to 50 °C. Solution 4 was 30 ml of filter-sterilized 10% (w/v) casamino acids. After 4h the colour of the plates was investigated. A change towards yellow/orange was considered positive for siderophore-production.

### Iron-depleted medium

To represses siderophore-independent growth of microorganisms, iron depleted RPMI was used. Therefore, two times concentrated RPMI media 1640 (LifeTechnologies, Burlington, ON, Canada) was reconstituted from powder in ddH_2_O, supplemented with 2% w/v CA and treated with (7% w/v) Chelex100 resin (Bio-Rad, Mississauga, ON, Canada) at 4°C overnight. After sterile filtration 20% of complement-inactivated horse serum (Sigma) and 20 µM EDDHA were added and the solution was heated up to 50°C. Separately a 3% w/v agarose solution was prepared autoclaved and cooled to 50°C. Both solutions were mixed and plates were poured with 20 ml for each plate. Plates containing horse serum are referred to as HS-RPMI.

### Reciprocal usage of siderophores

For the spot assay nasal strains were cultured in TSB or BM liquid medium at 37°C for up to 7d (see Table S3). Bacterial were harvested and washed two times with RPMI containing 10 μM EDDHA. Bacterial isolates suspected to consume SPs were adjusted to OD_600_=0.05 and plated on HS-RPMI by using a cotton swab. Siderophore-producers were adjusted to OD_600_=4.0 and spotted on top (5 µl) of the dried background strain. The plates were incubated for 37℃ for 24h-7days (see Table S3).

### Plate based co-cultures assay

Test strains were grown in TSB broth for 24h-48h (see Table S3). Bacterial were harvested, washed two times with RPMI containing 10 μM EDDHA. Strains was adjusted to 2×10^3^ CFU/ml from which 100 µl was plated on RPMI-HS plates. Bacteria were added individually or mixed at a multiplicity of infection (MOI) ratio of 1:1 and incubated for 24 h (3°C). Bacterial colonies were imaged using a LEICA M125 microscope (1.25x magnification) and the diameter of bacteria colonies was calculated using Image J.

### Liquid based co-cultures assay

Staphylococcal strains were grown overnight (20 h) in TSB at 37°C with agitation. Cells were harvested and washed two times with RPMI containing 1% CA and 10 μM EDDHA. OD_600_ was adjusted to 1. For the co-cultivation 5 µl of *S. auerus::sGFP* together with 5 µl of competing strain were mixed with 500 µl of RPMI+1% CA+10 µM EDDHA+100 µg Holo-Transferrin per well (final OD_600_ of 0.02) in a 48-well microtiter plate (Nunc, Thermo Scientific). OD_600_ and fluorescence intensity (Ext._480 nm_/Em._525 nm_) was measured every 30 min for 24 h in a Tecan Spark microplate reader at 37°C orbital and linear shaking.

### Whole Genome Sequencing

The genomes of *Corynebacterium simulans* 50MNs_SDm2, *Corynebacterium pseudodiphtheriticum* 90VAs_B3, *Corynebacterium hesseae* 10VPs_Sm8, *Bacillus cereus* 45MNs_B5, *Mammaliicoccus sciuri* 9VPs_Sm2 and *Citrobacter koseri* 44VAs_B2 sequenced using long and short read sequencing.

For Illumina short read sequencing, DNA was isolated from cell pellets using QIAGEN’s DNeasy® PowerSoil® Pro Kit according to the manual’s instructions with 2 minutes of vortexing in PowerBead Pro tubes. Libraries were prepared using the Illumina DNA Prep (M) Tagmentation kit according to the manufacturer’s protocol with 500 ng DNA input and 5 cycles indexing PCR. Libraries were checked for correct fragment length on an Agilent 2100 BioAnalyzer and pooled equimolarly. The pool was sequenced on an Illumina MiSeq® v3 600 cycles Flow Cell with 2 x 150 bp read length and a depth 70x genome coverage.

For long read sequencing, the cell pellet was resuspended in 600 µL ATL buffer (Qiagen #939011) and transferred to a ZR BashingBead Lysis Tube (Zymo Research #S6012-50). The tube was vortexed horizontally for 2 minutes on a vortex shaker. To optimize the DNA extraction the supernatant was taken off and digested with RNAse A (Qiagen). The DNA was then automatically purified with the QIAamp 96 QIAcube HT kit (Qiagen #51331) with additional proteinase K on a QIAcube HT following the manufacturer’s instructions. For library generation, the Ligation Sequencing Kit (LSK) 109 (Oxford Nanopore) was used with native barcoding, following the manufacturer’s protocol, with 500 ng DNA input per sample and prolongated incubation times. Library size was assessed on a FEMTO Pulse (Agilent) and libraries were pooled equimolarly before sequencing on a FLO-PRO002 flow cell on a Nanopore PromethION device. 14.5 million reads with 55 Gb were generated.

DNA sequence reads were assembled using nf-core pipeline bacass (v2.0.0 (67)). Assembled scaffolds were annotated using PGAP (NCBI, v.2021-11-29.build5742 (68)) and curated using NCBI Genome Workbench (v3.7.0 (69)).

### Phylogenetic comparison of nasal bacteria

The 16S rRNA locus was used for highlighting relatedness of the members of the nasal microbiome used in this study. The representatives for all species besides *C. hessae* were obtained from the NCBI Refseq or RiboGrove database (70). For *C. hesseae* the respective locus was obtained from the WGS presented in this study. The resulting sequences were aligned using Clustal Omega featured in the msa R package (71). A phylogenetic tree was constructed using the UPGMA method provided by the phangorn package (72).

### Animal models and Ethics statement

All animal experiments were conducted in strict accordance with the German regulations of the Gesellschaft für Versuchstierkunde/Society for Laboratory Animal Science (GV-SOLAS) and the European Health Law of the Federation of Laboratory Animal Science Associations (FELASA) in accordance with German laws after approval (protocol IMIT 1/15 for cotton rat colonization) by the local authorities (Regierungspräsidium Tübingen). All, animal and human studies were carried out at the University Hospital Tübingen and conformed to institutional animal care and use policies. No randomization or blinding was necessary for the animal colonization models, and no samples were excluded. Animal studies were performed with cotton rats of both genders, 8-12 weeks old, respectively.

### Cotton rat nasal colonisation

For the colonization, a spontaneous streptomycin-resistant mutant of *S. auerus* Newman wild type was selected on BM agar plates containing 250 μg ml^−1^ streptomycin. Into this *S. auerus* Newman^strepR^ the cassette *fhuC::erm* was introduced using phage transduction from the Nebraska transposon mutant library (strain NE406_SAUSA300_0633). Cotton rats were anaesthetized and instilled with either 1 × 10^7^ *S. aureus* Newman^strepR^ or 1 × 10^7^ S. aureus Newman^strepR^ *fhuC::erm*. Five days after bacterial instillation, the animals were euthanized, and noses were surgically removed. The noses were vortexed in 1 ml of 1 × PBS for 30 s. Samples were plated on agar plates containing 250 μg ml^−1^ streptomycin using an EddyJet 2W to determine the bacterial CFU. The plates were incubated for 2 days under aerobic conditions.

### Statistical analysis

Statistical analysis was performed by using GraphPad Prism (GraphPad Software, Inc., La Jolla, USA; version 9). Statistically significant differences were calculated by appropriate statistical methods as indicated. P values of ≤0.05 were considered significant.

## List of abbreviations

SP – Siderophore; SF-A – staphyloferrin A; SF-B, Staphyloferrin B; HS-RPMIHorseserum RPMI; CA - casamino acids; EDDHA - ethylenediamine-di(o-hydroxyphenylacetic acid)

## Declarations

### Ethics approval and consent to participate

*In vivo* studies were approved by the local authority (Regierungspräsidium Tübingen IMIT01-21G). <u>Consent for publication</u>

All authors have read the manuscript and endorsed publication. <u>Availability of data and material</u>

Sequence data that support the findings of this study have been deposited NCBI with the primary accession PRJNA1028639. Other data is provided within the manuscript or supplementary information files.

### Authors’ contributions

Y.Z. and A.B performed and evaluated the majority *in vitro* experiments, JJ. P. assembled genomes and performed bioinformatic analysis. D. B., B.TS., and B.K. performed *in vivo* experiments. LA.A. constructed *ΔfhuA* strains for *in vivo* experiments. S.H designed the study, acquired funding and wrote the manuscript.

## Acknowledgements

We thank Prof. Andreas Peschel for helpful discussion of the experimental approaches of this study. We thank Prof. Timothy J. Foster and Libera Lo Presti for critically reading and editing this manuscript. Further we thank Karsten Becker and Martine Pestel-Caron for providing diverse human nasal isolates. We thank the NCCT (NGS Competence Center Tübingen) for performing whole genome sequencing.

---

Fig. S1: Siderophore production by *S. aureus* JE2 and siderophore production by isogenic mutants. Bacterial isolates were spotted on BHI-EDDHA agar in 24 well plates and incubated for one week at 37°C. After incubation wells were overlaid with CAS-containing top agar. Color change to yellow indicates siderophore production and was assessed **4** hours after the overlay.

Fig. S2: Growth of S. aureus USA300 LAC::sGFP. **Growth curves in iron-limited medium:** *S. aureus* USA 300 LAC*::sGFP and S. aureus USA300 LAC wildtype* were inoculated to a optical density OD 0.01 and grown for 18 h in 500 µl iron-limited medium (1x RPMI, 1% casamino acid, 10 µM EDDHA, 100 µg holo-transferrin) at 37°C under constant shaking in the Tecan Spark ® 10M multimode microplate reader. Growth was monitored via optical density at 600 nm and fluorescence intensity of the GFP signal (Ext. 480 nm and Em. 525 nm). The dotted line indicates the media control. Mean and SD of three to eight experiments is shown. **A)** Shows a comparison of the OD_600_ growth curves obtained by the *S. aureus* USA300 LAC wildtype and *S. aureus* USA 300 LAC*::sGFP,* and **B)** Shows of *S. aureus* USA 300 LAC*::sGFP* curve generated through the fluorescent signal*. **C)*** Correlation of OD_600_ and GFP signal (2,5 h – 11 h of the growth) show a linear relationship (formula y=12.54+1284*x)

Fig. S3. Iron dependent growth of S. aureus Newman^strepR^ *fhuC::erm* **A)** Strains were grown in the presence of 20 µM FeSO_4_, 200 nM aerobactin, 9.1% spent medium of Δ*sfa* (source of SF-B), 5,7% spent medium of Δ*sbn* (source of SF-A) as a sole source of iron. 500 µl of cultures were inoculated to an OD_600_= 0,05 in 48 well plates and OD_600_ was measured after 20 h using an Epoch1 plate reader. Mean and SD of three experiments are shown. Statistical analysis was performed using one-way ANOVA (<0,0001) with subsequent multiple comparison.

