## Supplementary material for "Nasal commensals reduce *Staphylococcus aureus* proliferation by restricting siderophore availability": Supl. Figures S1-S3

***S. aureus* JE2 wildtype** | ***S. aureus* JE2 Δ*sbn*** | ***S. aureus* JE2 Δ*sfa*** | ***S. aureus* JE2 Δ*sbn*Δ*sfa***

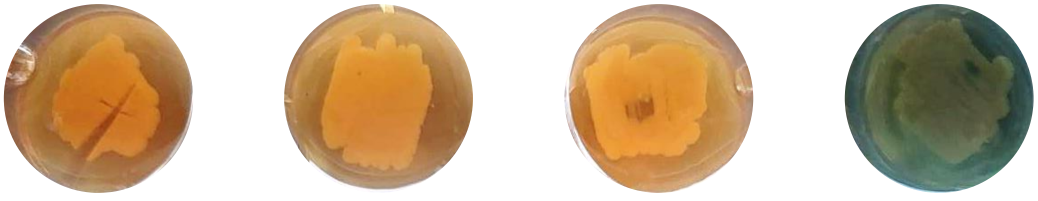

**Fig. S1: Siderophore production by *S. aureus* JE2 and siderophore production by isogenic mutants.** Bacterial isolates were spotted on BHI-EDDHA agar in 24 well plates and incubated for one week at 37°C**.** After incubation wells were overlaid with CAS-containing top agar. Color change to yellow indicates siderophore production and was assessed **4** hours after the overlay.

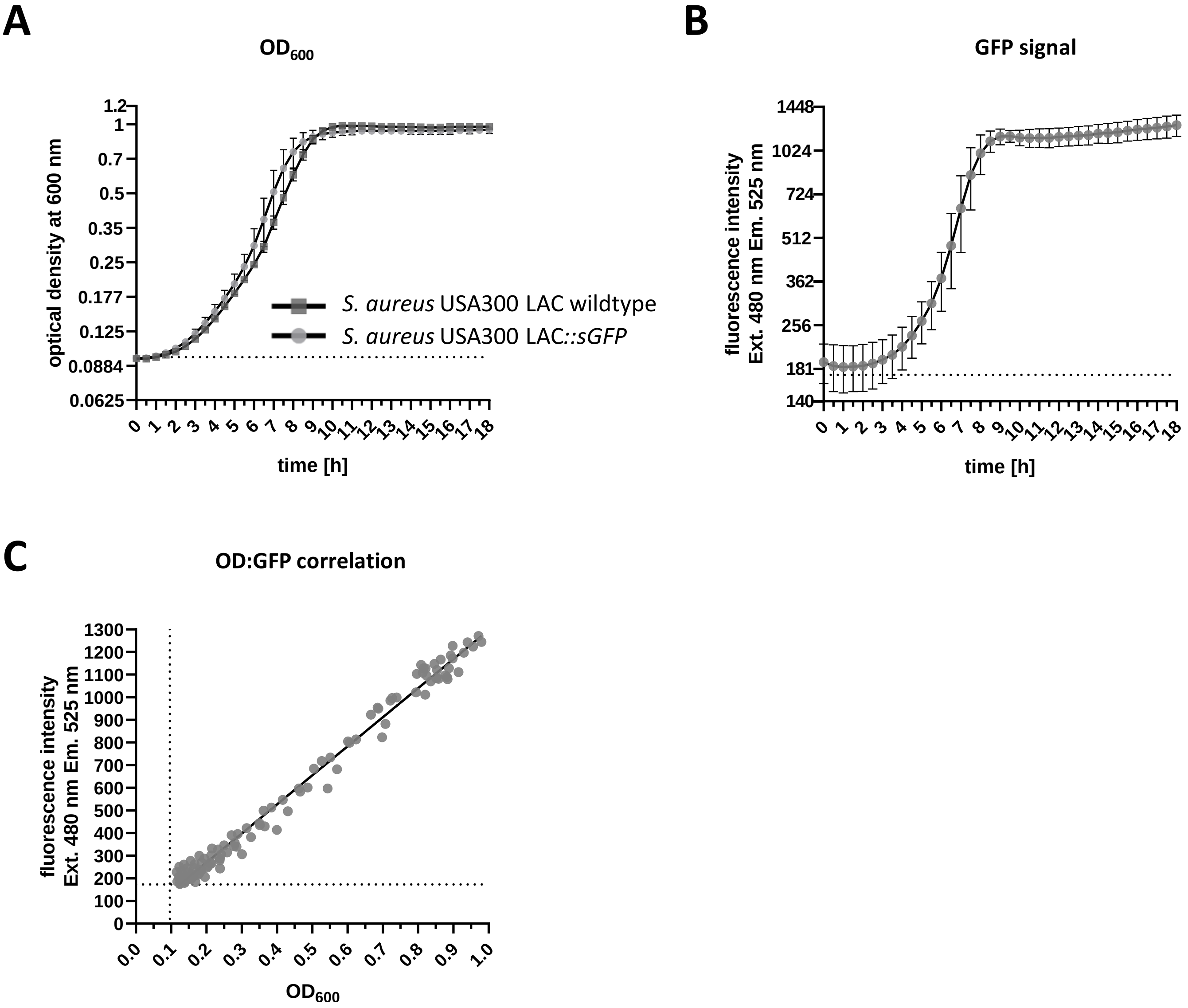


**Fig.S2: Growth of *S. aureus* USA300 LAC::*sGFP*.**

**Growth curves in iron-limited medium:** *S. aureus* USA 300 LAC*::sGFP and S. aureus* USA300 LAC wildtype were inoculated to a optical density OD 0.01 and grown for 18 h in 500 µl iron-limited medium (1x RPMI, 1% casamino acid, 10 µM EDDHA, 100 µg holo-transferrin) at 37°C under constant shaking in the Tecan Spark ® 10M multimode microplate reader. Growth was monitored via optical density at 600 nm and fluorescence intensity of the GFP signal (Ext. 480 nm and Em. 525 nm). The dotted line indicates the media control. Mean and SD of three to eight experiments is shown. **A)** Shows a comparison of the $OD_{600}$ growth curves obtained by the *S. aureus* USA300 LAC wildtype and *S. aureus* USA 300 LAC::*sGFP,* and **B)** Shows of *S. aureus* USA 300 LAC*::sGFP* curve generated through the fluorescent signal. ***C)*** Correlation of $OD_{600}$ and GFP signal (2,5 h – 11 h of the growth) show a linear relationship (formula y=12.54+1284*x)

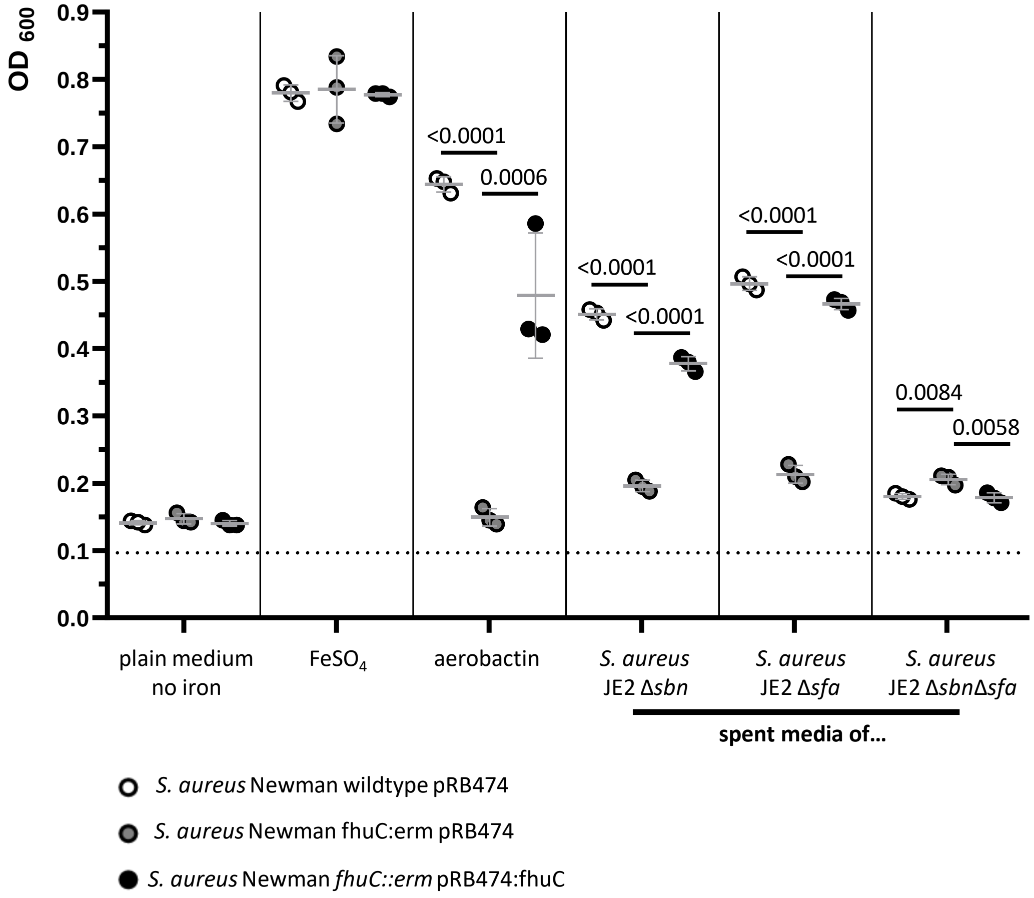


Fig. S3. Iron dependent growth of S. aureus Newman$^{strepR}$ *fhuC::erm.*

**A)** Strains were grown in the presence of 20 µM $FeSO_4$ , 200 nM aerobactin, 9.1% spent medium of Δ*sfa* (source of SF-B), 5,7% spent medium of Δ*sbn* (source of SF-A) as a sole source of iron. 500 µl of cultures were inoculated to an $OD_{600}$= 0,05 in 48 well plates and $OD_{600}$ was measured after 20 h using an Epoch1 plate reader. Mean and SD of three experiments are shown. Statistical analysis was performed using one-way ANOVA (<0,0001) with subsequent multiple comparison.
