## Supplementary material for "Nasal commensals reduce *Staphylococcus aureus* proliferation by restricting siderophore availability": Supl. Tables S1-S3

**Table S1. SirA and HtsA homologues in selected nasal commensals:**

| Species and strain designation | S. aureus receptor | Commensal Ortholog Protein ID | Identity (%) |
|---|---|---|---|
| *Corynebacterium hesseae* - 10VPs_Sm8 | HtsA | R3O64_11615 | 26 |
| | | R3O64_03755 | 26 |
| | SirA | R3O64_11615 | 34 |
| | | R3O64_03755 | 27 |
| *Citrobacter koseri* - 44VAs_B2 | HtsA | R3O63_20740 | 39 |
| | | R3O63_18180 (*fhuD*) | 28 |
| | SirA | R3O63_20740 | 23 |
| *Bacillus cereus* - 45MNs_B5 | HtsA | R3O67_00955 | 31 |
| | | R3O67_08985 | 33 |
| | | R3O67_21210 | 29 |
| | SirA | R3O67_00955 | 40 |
| | | R3O67_21210 | 37 |
| *Corynebacterium simulans* - 50MNs_SDm2 | HtsA | R3O68_02125 | 27 |
| | | R3O68_05280 (*fepB*) | 24 |
| | SirA | R3O68_05280 (*fepB*) | 28 |
| *Mammaliicoccus sciuri* - 9VPs_Sm2 | HtsA | R3O66_01910 | 46 |
| | SirA | R3O66_01220 | 40 |
| *Corynebacterium pseudodiphtheriticum* - 90VAs_B3 | HtsA | R3O65_07865 | 21 |
| | SirA | R3O65_07865 | 30 |
| | | R3O65_08540 | 62 |

**Table S2 Antismash-based prediction of biosynthetic gene clusters in selected nasal commensal isolates.**

| Species | Cluster | Function | Known Similarity | Identity % | Siderophore class |
|---|---|---|---|---|---|
| *Corynebacterium hesseae* 10VPs_Sm8 | 5.1 | T1PKS | | -- | |
| | 6.1 | NRPS-like | Dechlorocuracomycin | 8 | |
| | | | Desotamide | 9 | |
| | 11.1 | terpene | Carotenoid | 25 | |
| *Citrobacter koseri* 44VAs_B2 | 1.1 | siderophore | Aerobactin | 77 | Hydroxamate |
| | 2.1 | NRPS | Turnerbactin | 30 | Catecholate |
| | 5.1 | thiopeptide | O-antigen | 14 | |
| | 6.1 | NRPS, T1PKS | Yersiniabactin | 16/100 | Phenolate |
| | 7.1 | arylpolyene | APE Ec | 94 | |
| *Bacillus cereus* 45MNs_B5 | 2.1 | NRPS | | -- | |
| | 2.2 | NRPS | Bacillibactin | 100 | Catecholate |
| | 2.3 | siderophore | Petrobactin | 100 | Catecholate |
| | 3.1 | bacteriocin | | | |
| | | LAP | | | |
| | 5.1 | bateriocin | | | |
| | 6.1 | terpene | Molybdenum cofactor | 17 | |
| | 8.1 | bacteriocin | | | |
| | 12.1 | NRPS-like | | | |
| | 14.1 | betalactone | Fengycin | 40 | |
| | 14.2 | bacteriocin | | | |
| *Corynebacterium simulans* 50MNs_SDm2 | 3.1 | T1PKS | | -- | |
| | 3.2 | NRPS-like | Desotamide | 9 | |
| | 3.3 | NRPS | Glycopeptidolipid | 12 | |
| | 4.1 | bacteriocin | | | |
| | 6.1 | terpene | Carotenoid | | |
| *Mammaliicoccus sciuri* 9VPs_Sm2 | 2.1 | bacteriocin | | | |
| | 3.1 | terpene | | | |
| | 3.2 | siderophore | | | unkown |
| | 7.1 | lathipeptide | Nukacin ISK-1 | 18 | |
| *Corynebacterium pseudodiphtheriticum* 90VAs_B3 | 5.1 | terpene | | | |
| | 5.2 | T1PKS | | -- | |
| | 8.1 | NRPS | Cahuitamycins | 8 | |

**Table S3 List of human nasal isolates and their growth conditions**

| Species | Name | Source | Over-night cultures | | | Spott-Assay |
|---|---|---|---|---|---|---|
| | | | Cultivation | Media | Inc Time | Inc Time |
| *Bacillus cereus* | 44VPs_B5 | (22) | aerobic | TSB | 24 h | 1 d |
| | 45MNs_B5 | (22) | aerobic | TSB | 24 h | 1 d |
| | 89VPs_B11 | (22) | aerobic | TSB | 24 h | 1 d |
| *Bacillus mycoides* | 50MNt_Sm12 | (22) | aerobic | TSB | 24 h | 1 d |
| *Citrobacter koseri* | 10VAs_B1 | (22) | aerobic | TSB | 24 h | 1 d |
| | 44VAs_B2 | (22) | aerobic | TSB | 24 h | 1 d |
| | 9VAs_B2 | (22) | aerobic | TSB | 24 h | 1 d |
| *Corynebacterium accolens* | 10VAs_B6 | (22) | aerobic | BHI-Tween | 48 h | 2 d |
| | 50VPs_B7 | (22) | aerobic | BHI-Tween | 48 h | 2 d |
| | 63VAs_B8 | (22) | aerobic | BHI-Tween | 48 h | 2 d |
| | 83VAs_B5 | (22) | aerobic | BHI-Tween | 48 h | 2 d |
| *Corynebacterium aurimucosum* | 12UNs_B6 | (22) | aerobic | BHI-Tween | 48 h | 2 d |
| | 81VAs_KB1 | (22) | aerobic | TSB | 48 h | 2 d |
| *Corynebacterium hesseae* | 10VPs_Sm8 | (22) | aerobic | TSB | 24 h | 2 d |
| *Corynebacterium kroppenstedtii* | 82VAs_B6 | (22) | aerobic | BHI-Tween | 48 h | 3 d |

| | | | | | | |
|---|---|---|---|---|---|---|
| *Corynebacterium propinquum* | 63VAs_B4 | (22) | aerobic | BHI-Tween | 48 h | 2 d |
| | 80VAs_KB2b | (22) | aerobic | BHI-Tween | 48 h | 2 d |
| | 83VAs_B4 | (22) | aerobic | BHI-Tween | 48 h | 2 d |
| | 8VAs_B3 | (22) | aerobic | BHI-Tween | 48 h | 2 d |
| *Corynebacterium pseudo-diphtheriticum* | *M10-37* | (23) | aerobic | BHI-Tween | 48 h | 2 d |
| | *M8-43* | (23) | aerobic | BHI-Tween | 48 h | 2 d |
| | *P1.29* | (23) | aerobic | BHI-Tween | 48 h | 2 d |
| | *P2.34* | (23) | aerobic | BHI-Tween | 48 h | 2 d |
| | *P6. 3* | (23) | aerobic | BHI-Tween | 48 h | 2 d |
| | 44VPs_Sm3 | (22) | aerobic | BHI-Tween | 48 h | 2 d |
| | 87VAs_B4 | (22) | aerobic | BHI-Tween | 48 h | 2 d |
| | 90VAs_B3 | (22) | aerobic | BHI-Tween | 48 h | 2 d |
| *Corynebacterium simulans* | 50VAs_B5 | (22) | aerobic | TSB | 24 h | 1 d |
| | 50MNs_Sm2 | (22) | aerobic | TSB | 24 h | 1 d |

| | | | | | | |
|---|---|---|---|---|---|---|
| | 81MNs_B1 | (22) | aerobic | BHI-Tween | 48 h | 2 d |
| | 88UNs_Sm6 | (22) | aerobic | BHI-Tween | 48 h | 2 d |
| *Corynebacterium tuber-culostearicum* | 12VAs_B4 | (22) | aerobic | BHI-Tween | 48 h | 2 d |
| | 87VAs_B5 | (22) | aerobic | BHI-Tween | 48 h | 2 d |
| | 89VPs_B8 | (22) | aerobic | BHI-Tween | 48 h | 2 d |
| *Cutibacterium acnes* | 50VAs_Sa1 | (22) | anaerobic | TSB | 24 h | 1 d |
| | 83VAs_Sa3 | (22) | anaerobic | TSB | 24 h | 1 d |
| | 87VAs_SaT9 | (22) | anaerobic | TSB | 24 h | 1 d |
| | 89VAs_Sa2 | (22) | anaerobic | TSB | 24 h | 1 d |
| *Cutibacterium avidum* | 10VAs_Sa2 | (22) | anaerobic | TSB | 24 h | 1 d |
| | 63VAs_Sa1 | (22) | anaerobic | TSB | 24 h | 1 d |
| | 83VAs_Sa1 | (22) | anaerobic | TSB | 24 h | 1 d |
| | 89VAs_KBa2 | (22) | anaerobic | TSB | 24 h | 1 d |
| *Cutibacterium granulosum* | 50MNt_Sa3 | (22) | anaerobic | TSB | 24 h | 1 d |
| | 83VPs_KBa2 | (22) | anaerobic | TSB | 24 h | 1 d |
| *Dolosigranulum pigrum* | 9VAs_B4 | (22) | aerobic | TSB | 7 d | 7 d |
| | 90VAs_B6 | (22) | aerobic | TSB | 7 d | 7 d |
| *Finegoldia magna* | *87VAs_Sa4* | (22) | anaerobic | TSB | 4 d | 4 d |
| | 63VAs_Sa4 | (22) | anaerobic | TSB | 24 h | 1 d |
| | 83VAs_Sa6 | (22) | anaerobic | TSB | 48 h | 1 d |
| *Moraxella catarrhalis* | 44VAs_Sm4 | (22) | aerobic | TSB | 24 h | 1 d |
| | 80VAs_B4 | (22) | aerobic | TSB | 24 h | 1 d |

| | | | | | | |
|---|---|---|---|---|---|---|
| | 90VAs_B10 | (22) | aerobic | TSB | 24 h | 1 d |
| *Peptoniphilus harei* | 82VAs_KBa3 | (22) | anaerobic | TSB | 48 h | 3 d |
| *Staphylococcus aureus* | M11-28 | (23) | aerobic | TSB | 24 h | 1 d |
| | *M13-14* | (23) | aerobic | TSB | 24 h | 1 d |
| | *M15-5* | (23) | aerobic | TSB | 24 h | 1 d |
| *Staphylococcus capitis* | *M9-48* | (23) | aerobic | TSB | 24 h | 1 d |
| | *M11-5* | (23) | aerobic | TSB | 24 h | 1 d |
| | *M12-47* | (23) | aerobic | TSB | 24 h | 1 d |
| | 10VAs_KB2 | (22) | aerobic | TSB | 24 h | 1 d |
| | 44UNs_B2 | (22) | aerobic | TSB | 24 h | 1 d |
| | 50VAs_KB6 | (22) | aerobic | TSB | 24 h | 1 d |
| *Staphylococcus epidermidis* | *M11-13* | (23) | aerobic | TSB | 24 h | 1 d |
| | *M12-38* | (23) | aerobic | TSB | 24 h | 1 d |
| | *M14-1* | (23) | aerobic | TSB | 24 h | 1 d |
| *Staphylococcus hominis* | 50MNs_Sa6 | (22) | aerobic | TSB | 24 h | 1 d |
| | 89VPs_B7 | (22) | aerobic | TSB | 24 h | 1 d |
| | 9VPs_KB1 | (22) | aerobic | TSB | 24 h | 1 d |
| *Staphylococcus lugdunensis* | *SL2* | (73) | aerobic | TSB | 24 h | 1 d |
| | *SL9* | (73) | aerobic | TSB | 24 h | 1 d |
| | *SL13* | (73) | aerobic | TSB | 24 h | 1 d |
| | *SL27* | (73) | aerobic | TSB | 24 h | 1 d |
| | *SL37* | (73) | aerobic | TSB | 24 h | 1 d |
| | *SL57* | (73) | aerobic | TSB | 24 h | 1 d |
| | *SL62* | (73) | aerobic | TSB | 24 h | 1 d |
| | *SL71* | (73) | aerobic | TSB | 24 h | 1 d |
| | *SL72* | (73) | aerobic | TSB | 24 h | 1 d |

| | | | | | | |
|---|---|---|---|---|---|---|
| | *SL81* | (73) | aerobic | TSB | 24 h | 1 d |
| *Mammaliicoccus sciuri* | 9VPs_Sm2 | (22) | aerobic | TSB | 24 h | 1 d |
| *Staphylococcus warneri* | *M8-7* | (23) | aerobic | TSB | 24 h | 1 d |
| | *M11-4* | (23) | aerobic | TSB | 24 h | 1 d |
| | *M15-28* | (23) | aerobic | TSB | 24 h | 1 d |
| *Streptococcus constellatus* | *45UNs_KB5* | (22) | aerobic | TSB | 7 d | 7 d |
| *Streptococcus dysgalactiae* | 81VAs_B8a | (22) | aerobic | TSB | 48 h | 3 d |
| | 81VAs_KBa3a | (22) | aerobic | TSB | 48 h | 3 d |
| *Streptococcus intermedius* | 84MNs_KBa6 | (22) | aerobic | TSB | 48 h | 4 d |
| *Streptococcus mitis* | 78MNs_B4 | (22) | aerobic | TSB | 3 d | 4 d |
| | 81UNs_Sa2a | (22) | aerobic | TSB | 48 h | 4 d |
| *Streptococcus mutans* | 44UNt_B13 | (22) | aerobic | TSB | 48 h | 3 d |
| *Streptococcus oralis* | 44UNt_B6 | (22) | aerobic | TSB | 48 h | 3 d |
| *Streptococcus pneumoniae* | 80UNt_KB6 | (22) | aerobic | TSB | 48 h | 3 d |
